# Extensive NEUROG3 occupancy in the human pancreatic endocrine gene regulatory network

**DOI:** 10.1101/2021.04.14.439685

**Authors:** Valérie Schreiber, Reuben Mercier, Sara Jiménez, Tao Ye, Emmanuel García-Sánchez, Annabelle Klein, Aline Meunier, Sabitri Ghimire, Catherine Birck, Bernard Jost, Kristian Honnens de Lichtenberg, Christian Honoré, Palle Serup, Gérard Gradwohl

## Abstract

**Objective:** Mice lacking the bHLH transcription factor (TF) Neurog3 do not form pancreatic islet cells, including insulin secreting beta cells, causing diabetes. In human, homozygous mutations of *NEUROG3* manifest with neonatal or childhood diabetes. Despite this critical role in islet cell development, the precise function and downstream genetic programs regulated directly by NEUROG3 remain elusive. We therefore mapped genome-wide NEUROG3 occupancy in human induced pluripotent stem cell (iPSC)-derived endocrine progenitors and determined NEUROG3 dependency of associated genes to uncover direct targets.

**Methods:** We generated a novel hiPSC line (NEUROG3-HA-P2A-Venus), where NEUROG3 is HA-tagged and fused to a self-cleaving fluorescent VENUS reporter. We used the CUT&RUN technique to map NEUROG3 occupancy and epigenetic marks in pancreatic endocrine progenitors (PEP) differentiated from this hiPSC line. We integrated NEUROG3 occupancy data with chromatin status and gene expression in PEPs and their NEUROG3-dependence. In addition, we searched whether NEUROG3 binds type 2 diabetes mellitus (T2DM)-associated variants at the PEP stage.

**Results:** CUT&RUN revealed a total of 863 NEUROG3 binding sites assigned to 1268 unique genes. NEUROG3 occupancy was found at promoters as well as at distant cis-regulatory elements frequently overlapping within PEP active enhancers. *De novo* motif analyses defined a NEUROG3 consensus binding motif and suggested potential co-regulation of NEUROG3 target genes by FOXA, RFX or PBX transcription factors. Moreover, we found that 22% of the genes downregulated in *NEUROG3*^−/−^ hESC-derived PEPs are bound by NEUROG3 and thus likely to be directly regulated. NEUROG3 targets include transcription factors known to have important roles in islet cell development or function, such as *NEUROD1, PAX4, NKX2-2, SOX4, MLXIPL, LMX1B, RFX3*, and *NEUROG3* itself. Remarkably, we uncovered that NEUROG3 binds transcriptional regulator genes with enriched expression in human fetal pancreatic alpha (e.g., *IRX1, IRX2*), beta (e.g., *NKX6-1, SMAD9, ISX, TFCP2L1*) and delta cells (*ERBB4*) suggesting that NEUROG3 could control islets subtype programs. Moreover, NEUROG3 targets genes critical for insulin secretion in beta cells (e.g., GCK, ABCC8/KCNJ11, CACNA1A, CHGA, SCG2, SLC30A8 and PCSK1). In addition, we unveiled a panel of ncRNA potentially regulated by NEUROG3. Lastly, we identified several T2DM risk SNPs within NEUROG3 peaks suggesting a possible developmental role of NEUROG3 in T2DM susceptibility.

**Conclusion:** Mapping of NEUROG3 genome occupancy in PEPs uncovers an unexpectedly broad, direct control of the endocrine gene regulatory network (GRN) and raises novel hypotheses on how this master regulator controls islet and beta cell differentiation.

**Highlights:**

- NEUROG3 CUT&RUN analysis revealed 1268 target genes in human pancreatic endocrine progenitors (PEPs)
- NEUROG3 binding sites overlap with active chromatin regions in PEPs.
- 1/5 of the genes downregulated in *NEUROG3*^−/−^ hESC-derived PEPs are bound by NEUROG3.
- NEUROG3 targets islet specific TFs and regulators of insulin secretion.
- Several T2DM risk allelles lie within NEUROG3 bound regions.

## 1. INTRODUCTION

Diabetes results either from an auto-immune destruction of beta cells (Type 1 diabetes) or defective insulin secretion combined with the resistance of the peripheral tissues to insulin action (Type 2 diabetes). These forms of diabetes are considered as polygenic. On the other hand, mutations in single genes can also lead to rare early-onset forms of diabetes, thus defined as monogenic diabetes, for which the prevalence is estimated to be 2-5% of diabetes cases [1]. Monogenic diabetes is classified according to the age of onset and includes Neonatal Diabetes Mellitus (NDM) and Maturity Onset Diabetes of the Young (MODY), where diabetes occurs before 6 months and 25 years, respectively. These rare forms of diabetes result from mutations in genes controlling beta cell development, function, or both, including genes encoding essential transcription factors such as *PTF1A, PDX1, HNF1B, NEUROG3, RFX6*, or *NEUROD1* [1].

Among these genes, the bHLH transcription factor NEUROG3 is the key regulator of endocrine cell fate decision in the embryonic pancreas. In the mouse, all pancreatic islet cells derive from Neurog3-expressing pancreatic endocrine progenitors (PEP) and depend on *Neurog3* [2; 3]. *Neurog3*-deficient newborn mice die within a few days; they are diabetic since they lack insulin-secreting beta cells as well as all other islet cells [3]. In humans, homozygous or compound heterozygous mutations in *NEUROG3* have been identified in patients developing diabetes [4-7]. The pathology declared at various ages from neonatal to childhood, probably reflects differences in how severely NEUROG3 function is compromised. Of note, patients also developed rare forms of congenital malabsorptive diarrhea due to the lack of intestinal endocrine cells, which do not develop in the absence of NEUROG3 [4; 8]. Using pancreatic differentiation of human pluripotent stem cells as a model, it has been shown that *NEUROG3* is required for endocrine cell development [9; 10].

Despite its key function in endocrine commitment, the direct genetic program implemented by NEUROG3 is largely unknown both in mouse and human. Genome wide approaches have been performed to identify *Neurog3*-regulated genes in the mouse embryonic pancreas [11]. However, since the islet lineage is lost in the absence of Neurog3, a comparison of transcriptomes between Neurog3-deficient and control embryos revealed the entire islet transcriptome from endocrine progenitors to mature hormone-expressing cells, not only Neurog3 regulated genes. Direct Neurog3 target candidate genes such as *NeuroD, Nkx2-2, Insm1, Pax4, Neurog3* and *Cdkn1a* have been characterized previously using *in vitro* EMSA, Chromatin Immunoprecipitation (ChIP) and transactivation assays [12-16]. Using EMSAs and ChIP-qPCR, direct binding of NEUROG3 to *NKX2-2* and *NEUROG3* regulatory regions in hES-derived pancreatic precursors was recently reported [17]. Nevertheless, genome wide analysis to identify NEUROG3 bound regions and thus the entire panel of potential direct NEUROG3 targets has not been described. Such studies have been hampered by the lack of sensitivity of ChIP-Seq technique combined with the scarcity of NEUROG3-expressing endocrine progenitors.

Here we generated a novel hiPSC cell line where endocrine progenitor cells can be purified, and NEUROG3 is epitope tagged. We used the cleavage under targets and release using nuclease (CUT&RUN) technique, a method allowing transcription factor profiling from a low cell number [18-20], to identify NEUROG3 bound regions across the genome in hiPSC-derived pancreatic cells. We confirmed previously known NEUROG3 targets validating the experimental approach. Importantly, we identified many NEUROG3 targets that have not been reported before. Comparison with transcriptome data identified NEUROG3 bound genes that are enriched in human fetal pancreatic progenitors and regulated by NEUROG3. Our study uncovers an unexpectedly large panel of direct NEUROG3 targets in human pancreas progenitors, comprising an extensive endocrine GRN that implements NEUROG3 function.

## 2. MATERIALS AND METHODS

### 2.1. Culturing of iPSC lines

Wild type SB AD3.1 [21] and AD/N3HAV lines were maintained as undifferentiated hiPSC in mTeSR1 medium (Stem Cell Technology) on 1:30 diluted Matrigel (hESC grade, Corning) coated tissue culture surfaces, with everyday medium change. Cells were split every 3 or 4 days with TrypLE Select (Fisher) and seeded at 1.5-4×10e5 in a Matrigel-coated p35 plate containing 5μM Y27632 (Stem Cell Technologies) (mTeSR+Y) for the first day.

### 2.2. Generation of the NEUROG3-HA-P2A-Venus line

The SB AD3.1 line [21] was co-transfected with a pX458-plasmid expressing the sg1 guide RNA (Suppl. Table 1) and the Cas9 fused to GFP, and the targeting vector pBSII-KS-hNEUROG3-3HA-2A-3NLS-Venus-pA, both generated in the laboratory. Nucleofection was performed according to the manufacturer instructions (Amaxa), with 8×10e5 SB AD3.1 cells mixed with 2.5 μg each plasmid DNA, and cells were seeded on a p35 with mTeSR1+Y. The following day, cells were harvested with TryPLE, resuspended in PBS containing 2% FCS, 10 μM Y27632 and 1% Penicillin/Streptomycin, sorted by expression of GFP and seeded as 200 and 500 cells in several p35 in mTeSR1+Y. After 12 days, clones were picked by scratching and expanded for banking while genotyping.

### 2.3. Genotyping

DNA was purified from collected cells using the Nucleospin Tissue XS kit (Macherey-Nagel) according to the manufacturer instructions and genotyped by nested PCR using primers described in Suppl. Figure 1 and Suppl. Table 1. PCR products were purified using the Nucleospin Gel and PCR clean-up kit (Macherey-Nagel) and sequenced with appropriated primers (Suppl. Table 1) at Eurofins Genomics (Ebergberg, Germany).

### 2.4. Differentiation of hiPSC cells to pancreatic progenitors

Cells were differentiated according to the protocol of Petersen et al. (2017) [21]. At 80-90% confluency, cells were harvested with TryPLE and seeded at 3×10e5 cells/cm2 on Growth Factor Reduced Matrigel-coated 24wells- or 6wells-plates (CellBind Corning) in mTESR+Y. Differentiation was initiated 24 h after seeding. Cells were first rinsed with 1x PBS, then exposed daily to freshly prepared differentiation medium (Suppl. Table 1).

### 2.5. Flow cytometry analyses

Cells were harvested with TrypLE Select as described above, quenched with 3 volumes of MCDB131-3 medium containing 5 mM Y27632 (M3Y), washed once with PBS and fixed with 4% formaldehyde in PBS for 20 min. After 2 washes with PBS, cells were either stored at +4°C in PBS, BSA 1% for delayed analysis, or permeabilized 30 min with PBS, Triton 0.2%, 5% Donkey serum (permeabilization buffer) then incubated overnight at +4°C with primary antibodies (Suppl. Table 1) diluted in permeabilization buffer. After 2 washes with PBS-Triton 0.1%, 0.2% BSA (PBSTB), cells were incubated for 1-2 hour at RT with fluorophore-conjugated secondary antibodies (Suppl. Table 1) diluted in permeabilization buffer. After 2 washes with PBSTB, cells were resuspended at 1M/ml in PBS, 1% BSA, filtered on 85µm nylon mesh and analyzed on a BD Fortessa LSR II Cell analyser (BD Bioscience).

### 2.6. Immunofluorescence imaging

Cells were washed twice with PBS, fixed with 4% formaldehyde in PBS for 20 min, permeabilized 30 min with PBS-Triton 0.5% and blocked for 30 min in PBSTB. Cells were incubated with primary antibodies (Suppl. Table 1) diluted in PBSTB overnight at 4°C, washed 3x in PBS-Triton 0.1% and incubated for 1-2 hour at RT with fluorophore-conjugated secondary antibodies (Suppl. Table 1) diluted in PBSTB. Cells were washed twice in PBSTB, nuclei were stained with Dapi 50 ng/mL in PBST. Image acquisition was done on an inverted fluorescence microscope Leica DMIRE2.

### 2.7. Flow cytometry sorting of Venus+ cells

Cells were harvested with TrypLE Select at day 13 of differentiation, quenched with 3 volumes of M3Y, centrifuged 4 min at 200g, resuspended at 5M/ml in M3Y and sorted using a FACS Fusion/Aria in M3Y at +4°C. Venus+ cells were collected and either used immediately or cryoconserved in Cryostor10 (Stem Cell Technologies) at −80°C.

### 2.8. CUT&RUN

We followed the protocol of Hainer and Fazzio (2019) [22] with minor modifications. Freshly sorted cells (75,000 for anti NEUROG3, HA and CTRL donkey anti sheep (DAsh) antibodies and 18,000 cells for H3K4me3 antibody) or thawed sorted cells (15,000 for H3K27me3 and rabbit anti mouse control antibodies) were washed once with 1mL cold PBS and resuspended in nuclear extraction buffer (NE, 20mM HEPES-KOH, pH 7.9, 10mM KCl, 0.5mM Spermidine, 0.1% Triton X-100, 20% glycerol, freshly added protease inhibitors). After 3 min spinning at 4°C at 600g, cells were resuspended in 600 μL NE buffer. Concanavalin A beads (Polysciences, 25 μL bead slurry/sample) were washed twice with ice-cold Binding buffer (20mM HEPES-KOH, pH 7.9, 10mM KCl, 1mM CaCl2, 1mM MnCl_2_) and resuspended in 300 μL Binding buffer. Nuclei were added to beads with gentle vortexing and incubated for 10 min at 4°C with gentle rocking. Bead-bound nuclei were blocked with 1 mL cold Blocking buffer (20mM HEPES, pH 7.5, 150mM NaCl, 0.5mM Spermidine, 0.1% BSA, 2mM EDTA, freshly added protease inhibitors) by gentle pipetting, incubated 5 min at RT and washed in 1mL cold Wash buffer (WB, 20mM HEPES, pH 7.5, 150mM NaCl, 0.5mM Spermidine, 0.1% BSA, freshly added protease inhibitors) and resuspended in 250 μL cold WB. 250 μL of primary antibody (Suppl. Table 1) diluted 1:100 in cold WB were added with gentle vortexing, and samples were incubated overnight with gentle rocking at 4°C. Samples were washed twice in 1 mL cold WB, and resuspended in 250 μL cold WB. When indicated, incubation with a secondary antibody (Donkey anti Sheep IgG, 1:200) was performed for 1h at 4°C in WB under gentle rocking. After 2 washes with 1 ml WB, and resuspension in 250 μl WB, 200 μL of pA-MN (diluted at 1.4 ng/mL in cold WB) was added with gentle vortexing, and samples were incubated with rotation at 4°C for 1 hour. The pA-MN was produced in-house, according to the protocol described by [23] and using the pK19pA-MN plasmid, obtained from Ulrich Laemmli (RRID:Addgene_86973; http://n2t.net/addgene:86973). Samples were washed twice in 1 mL cold WB and resuspended in 150 μL cold WB. Three μL of 100 mM CaCl_2_ were added upon gentle vortexing to activate the MN. After 30 min of digestion, reactions were stopped by addition of 150 μL 2XSTOP buffer (200mM NaCl, 20mM EDTA, 4mM EGTA, 50ug/mL RNaseA, 40ug/mL glycogen) and DNA fragments were released by passive diffusion during incubation at 37°C for 20 min. After centrifugation for 5 min at 16,000g at +4°C to pellet cells and beads, 3 μL 10% SDS and 2.5 μL Proteinase K 20mg/ml were added to the supernatents, and samples were incubated 10 min at 70°C. DNA purification was done with phenol/chloroform/isoamyl alcohol extraction followed by chloroform extraction using MaxTract tubes (Qiagen). DNA was precipitated with ethanol after addition of 20 µg glycogene, and resuspended in 36.5µl 0.1XTE.

### 2.9. High throughput sequencing

Illumina sequencing libraries were prepared at the Genomeast facility (IGBMC, Illkirch). CUT&RUN samples were purified using Agencourt SPRIselect beads (Beckman-Coulter). Libraries were prepared from 10 ng of double-stranded purified DNA using the MicroPlex Library Preparation kit v2 (Diagenode) following the manufacturer’s protocol with some modifications. Illumina compatible indexes were added through a PCR amplification (3 min at 72°C, 2 min at 85°C, 2 min at 98°C; [20 sec at 98°C, 10 sec at 60°C] x 13 cycles). Amplified libraries were purified and size-selected using Agencourt SPRIselect beads (Beckman Coulter) by applying the following ratio: volume of beads / volume of libraries = 1,4 / 1. The libraries were sequenced on Hiseq 4000 as Paired-End 2×100 base reads following Illumina’s instructions.

### 2.10. Bioinformatics analyses

#### 2.10.1. Data processing

Image analysis and base calling were performed using RTA 2.7.3 and bcl2fastq 2.17.1.14. Reads were trimmed using cutadapt v1.9.1 with option: -a AGATCGGAAGAGCACACGTCTGAACTCCAGTCAC -A AGATCGGAAGAGCGTCGTGTAGGGAAAGAGTGTA -m 5 -e 0.1. Paired-end reads were mapped to Homo Sapiens genome (assembly hg38) using Bowtie2 (release 2.3.4.3, parameter: -N 1 -X 1000). Reads overlapping with ENCODE hg38 blacklisted region V2 were removed using Bedtools. Reads were size selected to <120bp and >150bp, since it has been reported that small reads define more precisely TF binding site, whereas larger reads (>150bp) result from sites occupied by nucleosomes [18; 19]. Bigwig tracks were generated using bamCoverage from deepTools for ≤120 bp and ≥150 bp fragments separately. Tracks were normalized with RPKM method. The bin size is 20. ≤120 bp fragments are used for samples obtained with anti NEUROG3 (VLSR28), HA (VLSR27) and the control donkey anti sheep (DAsh, thereafter named CTRL, VLSR29) antibodies and ≥150 bp fragments for samples obtained with anti H3K4me3 (VLSR32), anti H3K27me3 (VLSR44) and the rabbit anti mouse control (RAM, VLSR41) antibodies. Bigwig tracks (reads <120 bp long for NEUROG3, HA and CTRL samples and >150 pb for histone marks) were displayed on the reference genome *hg38* using UCSC genome browser. For simplicity, only the DAsh CTRL is illustrated throughout the manuscript. Heatmaps and K-means clustering was done using seqMINER v1.3.3g [24]. To compare with previously published data obtained from human *in vitro* derived pancreatic endocrine progenitors [25], multipotent progenitors [26] and from adult islets [27], we converted coordinates of bed and bigwig files to *hg19* coordinates using the UCSC Liftover and the bigwigLiftOver tools (https://github.com/milospjanic/bigWigLiftOver), respectively. Genomic tracks were visualized using http://meltonlab.rc.fas.harvard.edu/data/UCSC/SCbetaCellDiff_ATAC_H3K4me1_H3K27ac_WGBS_tracks.txt.

#### 2.10.2. Peak calling

Peak calling was performed with the Sparse Enrichment Analysis for CUT&RUN SEACRv1.3 tool [28] (https://seacr.fredhutch.org), using the norm and stringent mode on the <120bp size selected reads and VLSR29 (DAsh CTRL) as a control for VLSR27 (HA) and VLSR28 (NEUROG3) datasets. To identify overlapping genomic regions, peak coordinates were intersected using the BEDtools 2.22.0 command *intersect interval files* (http://use.galaxeast.fr).

#### 2.10.3. Association of peaks with genomic features and genes

Genomic annotation was first performed using the HOMER v3.4 [29] *annotatePeaks*.*pl* script with the default settings (promoters-transcription start site (TSS) from –1 kb to +100 bp to the TSS and transcription termination sites (TTS) from –100 bp to +1 kb of the TTS. GREAT 4.0.4 [30] was used to assign NEUROG3/HA peaks to their nearest coding gene(s) using basal settings (each gene is assigned a basal regulatory domain of 5 kb upstream and 1 kb downstream of its TSS. The gene regulatory domain is extended in both directions to the nearest gene’s basal domain but no more than 1,000 kb extension in one direction. Each peak is associated with all genes whose regulatory domain it overlaps). The NEUROG3 peaks or the distal peaks defined by GREAT (>5kb from TSS) were intersected with enhancers regions of hESC-derived endocrine progenitors (EN) lifted over to the *hg38* genome ([25], GSE139816).

#### 2.10.4. Motifs identification and analyses

*De novo* motif discovery and known motifs enrichment analysis were performed using the HOMER v3.4 [29] *findMotifsGenome*.*pl* script with default settings (200-bp windows centred on peak summits, motif lengths set to 8, 10 and 12 bp, hypergeometric scoring). For the 6 most significant *de novo* motifs identified, known best match motifs were associated if their Homer score was >0.85. Known co-occurring motifs were manually curated to exclude redundant bHLH motifs. Co-occurance of the *de novo* identified NEUROG3 motif and known RFX6 or FOXA2 motifs was done on the entire peak sequences using the HOMER script *annotatePeaks*.*pl* with -size given and -m <motif*n*.motif> options.

#### 2.10.5. Functional annotations

Gene functional annotation and clustering was carried out with DAVID v6.8 (https://david.ncifcrf.gov/home.jsp, [31]), using GO Biological Process, GO Cellular Component and KEGG Pathways. Selected terms significantly enriched and sorted by -Log(P-value) are displayed. To identify NEUROG3 transcription factors target genes, the peaks-assigned genes names were intersected by Venny 2.1.0 (https://bioinfogp.cnb.csic.es/tools/venny/) with a list of 1734 TF combining the 1639 human TF identified by [32] with the 1496 human TF taken from the human protein atlas (https://www.proteinatlas.org) (Suppl. Table 2). To identify ncRNA genes at the vicinity of NEUROG3 binding sites, we extended the genomic coordinates of the 2226 *de novo* and 2112 previously annotated LncRNA listed by Akerman et al (2017) [33] by 100kb (or 5kb) in both directions and intersected with *hg19* converted coordinates of NEUROG3 peaks.

#### 2.10.6. Overlap between bound genes and differentially expressed genes

Differential expression analysis between *NEUROG3*^−/−^ hESC line differentiated to day 13 and its wild-type counterpart, from corresponding RNA-seq data [34], was performed using a negative binomial GLM fit and Wald significance test implemented in the Bioconductor package DESeq2 version 1.16.1 [35]. The variables considered for the GLM model were the batch and condition. Differentially expressed genes were defined as those having a Benjamini – Hochberg-adjusted Wald test P < 0.05. A total of 319 genes were differentially expressed, from which 312 were downregulated in *NEUROG3*^−/−^ cells. The list of genes significantly enriched in NEUROG3-eGFP^+^ human pancreatic and endocrine progenitors differentiated *in vitro* from hESC cells (2852 genes), compared to NEUROG3-eGFP^−^ cells, was taken from Liu et al, 2014 [36]. Both gene lists were intersected with NEUROG3 bound genes list by Venny 2.1.0. Expression of genes of interest in human fetal pancreas and during *in vitro* differentiation of human embryonic stem cells to pancreatic endocrine cells was examined using https://descartes.brotmanbaty.org [37] and http://hiview.case.edu/public/BetaCellHub/differentiation.php [38], respectively.

#### 2.10.8. Enrichment of T2D-FG associated variants

The NEUROG3 bound EN_enhancers (hg19) and the NEUROG3-bound regions (hg19) were intersected using Bedtools 2.22.0 with the 23,144 genetic variants associated with T2D and glycemic traits (T2D-FG) on 109 loci, compiled by Miguel-Escalada et al [27].

#### 2.10.9. Data availability

Raw data have been deposited in the GEO database under accession code GSE171963. hESC-derived *NEUROG3*^−/−^ [34] and hESC-derived NEUROG3-eGFP^+^ cell [36] RNA-seq data are from E-MTAB-7185 and GSE54879, respectively. hESC-derived endocrine progenitors (EN) data (enhancers, H3K27ac ChIP-seq and RNA-seq, Ref [25]) are from GSE139817.

## 3. RESULTS AND DISCUSSION

### 3.1 Identification of NEUROG3 targets in hiPSC-derived endocrine progenitors

To unveil the endocrinogenic program implemented by NEUROG3, we mapped NEUROG3 occupancy across the genome during directed differentiation of hiPSC into beta cells. We first generated an hiPSC line where NEUROG3 is tagged with 3 HA epitopes and fused to a cleavable nuclear VENUS fluorescent reporter (NEUROG3-HA-P2A-Venus) (Figure 1A and Suppl. Figure 1). Using the protocol described by Petersen et al (2017) [21] and adapted from Rezania et al. (2014) [39], we differentiated the NEUROG3-HA-P2A-VENUS hiPS cells along the pancreatic and islet lineage until the pancreatic endocrine progenitor (PEP) stage 5, at day 13 (Figure 1A). By immunofluorescence, we found that NEUROG3-positive cells are indeed co-expressing HA and Venus, as expected (Suppl. Figure 2A-B). Accordingly, FACS analyses showed a correlation between HA and Venus expression (Suppl. Figure 2C). All the Venus+ cells are expressing PDX1 (Suppl. Figure 2A,C), as expected and previously shown with a NEUROG3-eGFP hiPSC line [21]. To map NEUROG3-bound regions, we used the CUT&RUN technique, an alternative to ChIP-seq for low input cell numbers [18; 19]. This technique is based on the recruitment of micrococcal nuclease, fused to protein A (pA-MNase), to antibody-bound sites within the genome in intact nuclei (Figure 1A). The subsequently cleaved fragments are recovered and sequenced. Endocrine progenitors were purified at day 13 (d13) of differentiation (Suppl. Figure 2D), and CUT&RUN experiments were performed on Venus+ cells using anti-NEUROG3 and anti-HA antibodies. To map chromatin states, we also profiled active (H3K4me3) and repressive (H3K27me3) histone marks.

**Figure 1.**
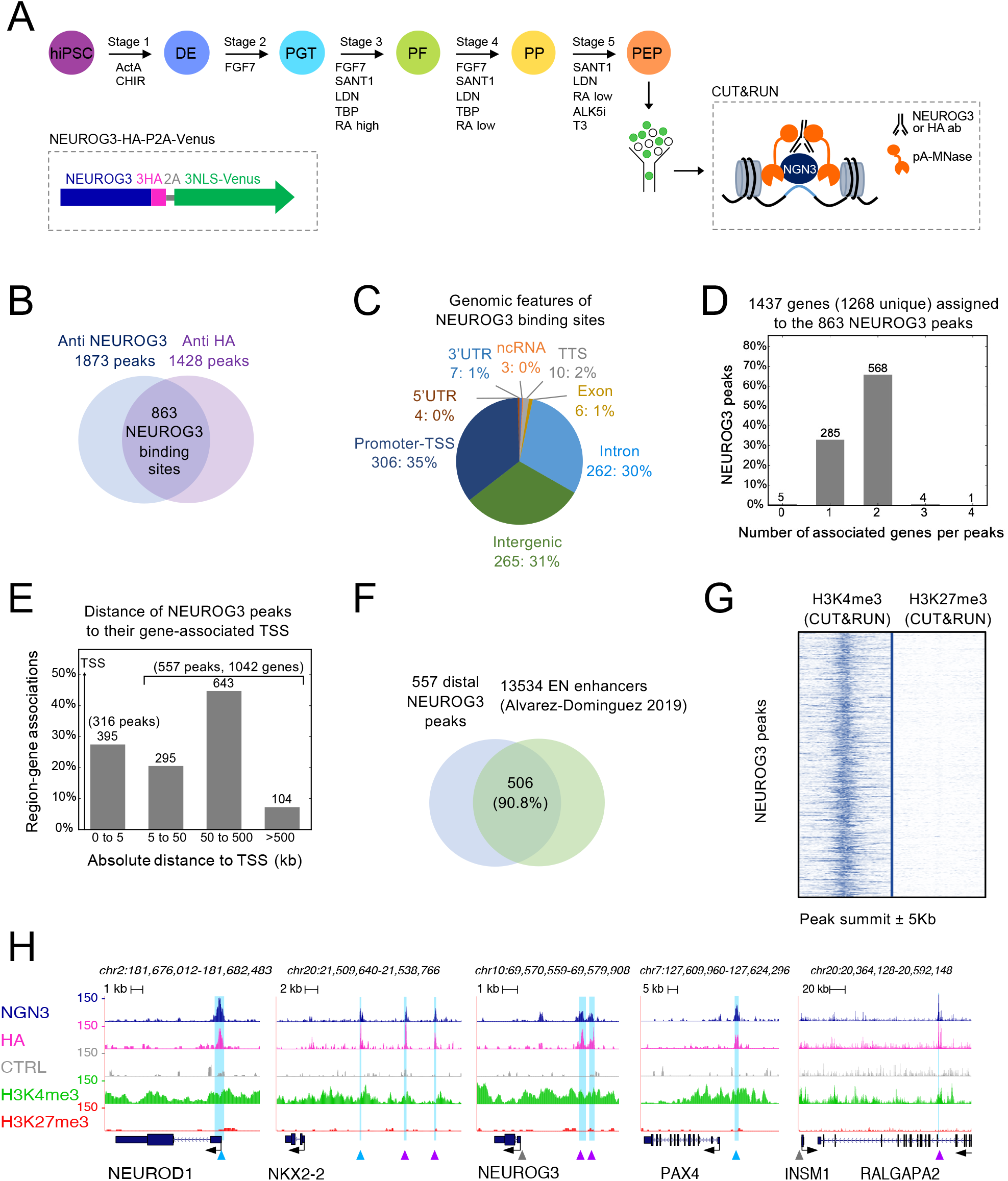
Characterization of the genome-wide binding sites of NEUROG3 in human hiPSC-derived pancreatic endocrine progenitors. (A) Overview of the study: a 5-stage protocol was used to differentiate hiPSC to pancreatic endocrine progenitors using the sequential supplementation of factors indicated. At day 13, Venus+ cells were sorted and used in a CUT&RUN experiment. Inset: schematic representation of the NEUROG3-3HA-P2A-3NLS-Venus allele. (B) Venn diagram showing the number and overlap of peaks identified by CUT&RUN with an anti-NEUROG3 or an anti-HA antibody. (C) Genomic distribution (number and % of peaks) of the 863-high confidence NEUROG3 binding sites. (D) Number of genes assigned by GREAT per NEUROG3 peaks. (E) Distance of NEUROG3 peaks to their gene(s)-associated TSS. (F) Overlap between NEUROG3 distal binding sites (>5kb from TSS) and enhancers regions of hiPSC-derived endocrine progenitors (EN), as defined by [25]. (G) Normalized read density surrounding NEUROG3 peak summit ±5Kb for H3K4me3 and H3K27me3 CUT&RUN data sets. (H) Genome browser tracks showing NEUROG3, HA, H3K4me3, H3K27me3 and the CTRL (Donkey anti Sheep antibody) CUT&RUN data at the *NEUROD1, NKX2-2, NEUROG3, PAX4* and *INSM1*/*RALGAPA2* loci. Identified NEUROG3 peaks are highlighted in light blue. Peaks matching previously reported NEUROG3 binding sites are indicated by blue arrowheads, newly discovered peaks by purple arrowheads, and reported sites not confirmed here by grey arrowheads [12; 13; 15-17].

We identified 1873 and 1428 peaks using NEUROG3 and HA antibodies, respectively (Figure 1B). To enhance the stringency of NEUROG3-bound regions, we intersected both datasets, defining NEUROG3 occupancy at 863 common sites (Figure 1B and Suppl. Table 2). These high confidence NEUROG3 binding sites were found at promoters (35%), in introns (30%), and in intergenic regions (31%) (Figure 1C) and were assigned by GREAT to 1268 unique genes, with 573 peaks (66%) assigned to 2 or more genes (Figure 1D). NEUROG3 binding to distal regions (located >5kb from the TSS of their associated gene) was observed for 65% of sites (557 peaks for 1042 genes) (Figure 1E). Remarkably, 90.8% (506 peaks) of these distal NEUROG3 bound regions were located within enhancer regions of hESC-derived endocrine progenitors (EN), as defined through their H3K27 acetylation by Alvarez-Dominguez et al. (2019) [25] (Figure 1F). In agreement, we found that H3K4me3 active histone marks were enriched at the NEUROG3 peaks compared to the H3K27me3 repressive marks (Figure 1G), indicating NEUROG3 binding at active promoters and enhancers. Taken together, we uncovered the NEUROG3 cistrome in PEPs, suggesting that NEUROG3 activates gene transcription by binding both proximal and distal cis-regulatory elements.

### 3.2 CUT&RUN detects previously identified and novel binding sites in known NEUROG3 targets

To validate the CUT&RUN approach for identifying of NEUROG3 bound regions in PEPs, we first examined previously characterized direct targets. As expected, peaks were identified in *NEUROD1, NKX2-2, PAX4, INSM1*, and *NEUROG3* [12; 13; 15-17], some of which at sites already mapped by ChIP-qPCR and/or EMSA and luciferase assays (Figure 1H). Interestingly, we identified two unreported NEUROG3 binding sites upstream of *NKX2-2* gene and upstream of *NEUROG3* TSS (purple arrowheads in Figure 1H). The sites identified for *NEUROG3* were distinct from the one reported previously by ChIP-qPCR [17] but overlapped with the conserved *Neurog3* enhancer region described in the mouse [40], supporting that NEUROG3 regulates its own transcription [12]. The peak assigned to *INSM1* is distantly located >180kb downstream of its TSS, within an intron of the *RALGAPA2* genes. However, this region has been identified as a super-enhancer directly linked to the *INSM1* gene in promoter capture HiC studies performed in adult pancreatic islet [27] (Figure 1H and Suppl. Figure 3) suggesting a function in the regulation of *INSM1* expression. Of note, we found no binding site for the *CDKN1A* gene, shown in the mouse to be directly regulated by NEUROG3 and to promote cell cycle exit in PEP [14]. It is possible that the NEUROG3 target NEUROD1 serves as an intermediate since NEUROD1 was shown to similarly inhibit cell proliferation by directly regulating *Cdkn1a* transcription [41]. Altogether, these data validate the CUT&RUN technique to unravel NEUROG3 bound sites genome-wide and suggest that the expected NEUROG3-driven endocrinogenic programs are activated in hiPSC-derived PEP.

### 3.3 Consensus NEUROG3 binding motif and co-binding of transcription factors

To determine the motifs enriched in the NEUROG3 binding regions, we performed a *de novo* motifs analysis [29] that revealed a strong enrichment for the RCCATCTGBY E-box type motif (CANNTG) recognized by bHLH transcription factors (Figure 2A). The NEUROG3 recognition motif is similar to NEUROD1 and NEUROG2 binding motifs, in agreement with the strong homology of the bHLH DNA binding domains between NEUROD and NEUROG families. Several additional motifs were found significantly enriched, such as the motif recognized by NFY, FOX, SP/KLF, RFX, and PBX TFs (Figure 2A-C and Suppl. Figure 4). Some TFs of these families have been reported to regulate pancreas development and islet cell differentiation, such as Pbx1 [42], Rfx3 and Rfx6 [43; 44]. Interestingly, the binding of the general NFY factors was reported biased towards regulatory elements with enhancer activity [45]. In agreement with our findings, KLF, FOXA1/A2, RFX, and MEIS1 (a PBX1 related homeobox gene) TFs have recently been predicted to bind to PEP Super Enhancers in a model of Core transcriptional regulatory circuits (CRCs) in the human islet lineage [25]. Of particular interest are the presence of FOX and RFX motifs in NEUROG3 bound regions. Indeed, FOXA1 and FOXA2 have been shown to act as pioneer factors facilitating chromatin access to other TFs at multiple stages during pancreas development [46]. 28.27% of the NEUROG3 peaks harbor a FOXA2 motif (Figure 2B). Moreover, 189 (21.97%) of NEUROG3 binding sites are bound by FOXA2 in *in vitro*-derived pancreatic multipotent progenitor cells (MPC) [26], and 36 of these sites match with *cis*-regulatory modules (CRM), binding sites of multiple TFs essential for early pancreas development, among which 15 are regulator elements of TF genes (Figure 2D and 2E, and see below) [26]. The pioneer activity of FOXA2 that was also described during human *in vitro* pancreatic progenitor differentiation [47] could therefore be required for the subsequent gene activation mediated by NEUROG3 at primed enhancers. The fact that FOXA2 possibly regulates *NEUROG3* (as shown in mouse [40]) and our findings that NEUROG3 binds *FOXA2* (Figure 2E) provides a possible additional regulatory loop between these two TFs. Interestingly, we identified a RFX6 motif in 37.54% of NEUROG3 peaks (Figure 2B) and revealed the co-occurence of the NEUROG3 motif with the RFX6 motif in 1/5 of the peaks, from which 1/3 had an additionnal FOXA2 motif (Figure 2C). Several NEUROG3 bound genes were indeed previously identified as Rfx6 targets in a mouse beta cell line [43] (Figure 2C and data not shown). Altogether, FOXA2, as well as RFX6, might be important coregulators of the transcription of NEUROG3 direct targets.

**Figure 2.**
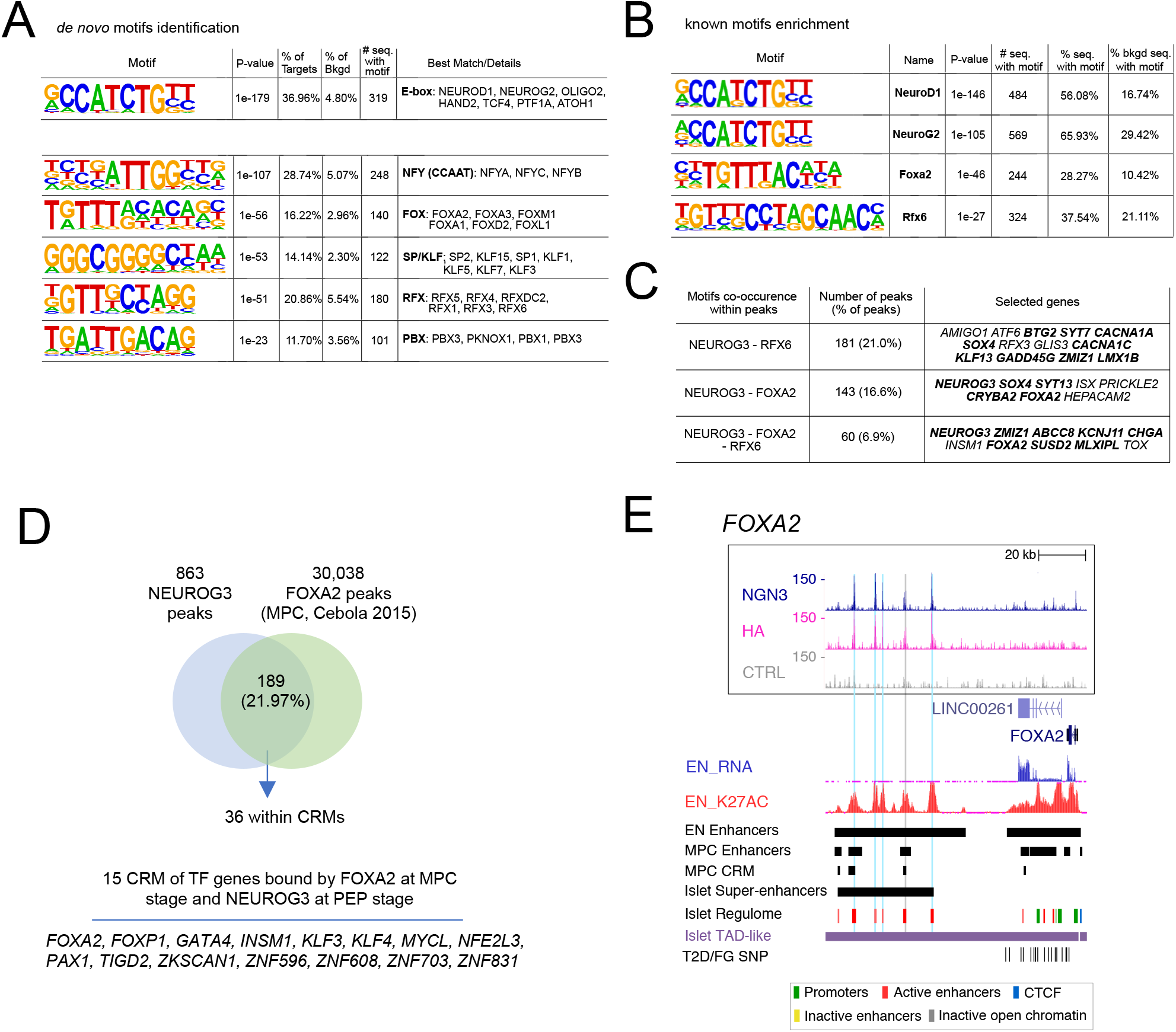
TF motifs discovery in NEUROG3 binding sites. (A). *De novo* motif discovery ranked by P-value reflecting motif enrichment within peak summit ±100bp. Number (#) and % of target and background sequences harboring each motif is indicated for the 6 most significant motifs identified. Known best matching transcription factors were associated if their HOMER score was >0.85. (B) Selection of known co-occuring motifs ranked by P-value within the entire peak sequences. The full list of the 50 most significantly enriched motifs is shown in Suppl Figure 4. (C) Co-occurence of NEUROG3 *de novo* identified motifs with motifs for RFX6 and/or FOXA2 on the entire peak sequences. Some selected GREAT assigned genes are indicated, and in bold when identified as targets for Rfx6 in mouse beta cell line [43]. (D-E) Regions bound by NEUROG3 at PEP stage that are bound by FOXA2 at the MPC stage. 36 regions coincide with MPC *cis*-regulatory modules (CRM) defined by [26], from which, 15 regulate TF genes (D). (E) Genome browser tracks showing NEUROG3, HA, and the CTRL CUT&RUN data at the *FOXA2* loci. Coordinates are from *hg19*. The position of NEUROG3 binding sites is highlighted in light blue (or in grey, when identified in a single dataset). Data of RNA-seq (EN_RNA), H3K27ac ChIP-seq (EN_K27AC) and the position of enhancers (EN Enhancers) from hESC-differentiated to endocrine progenitors were taken from [25] and http://meltonlab.rc.fas.harvard.edu/data/UCSC/SCbetaCellDiff_ATAC_H3K4me1_H3K27ac_WGBS_tracks.txt. Position of hESC-derived multipotent progenitor cells (MPC) enhancers and *cis*-regulatory modules (CRM, defined as regions bound by at least two TF) are taken from [26]. Data from adult islets (Super-enhancers, Islet regulome, T2D/FG SNP and TAD-like regions) are taken from [27], isletregulome.org and http://epigenomegateway.wustl.edu/.

### 3.4 Integration of NEUROG3 occupancy and gene expression in the islet lineage

Gene ontology (GO) analyses revealed that NEUROG3-bound regions are associated with genes retaled to GO terms such as endocrine pancreas development and insulin secretion, in agreement with the expected proendocrine function of NEUROG3 (Figure 3A and Suppl.Table 2). We, therefore, scrutinized NEUROG3 bound genes that are expressed in the islet lineage, expecting these genes to be downregulated in *NEUROG3*^−/−^ cells, or upregulated in NEUROG3-enriched cells. We thus used RNA-seq data comparing the transcriptome of *NEUROG3*^−/−^ versus wild-type hESC line, differentiated to d13 [34] and a previously published list of genes significantly enriched in NEUROG3-eGFP^+^ hESC-derived endocrine progenitors [36]. From the 319 differentially expressed genes in *NEUROG3*^−/−^ cells, 312 were downregulated (Suppl. Table 2) from which 69 (22%) were directly bound by NEUROG3 (Figure 3B-C, Suppl. Table 2). From the 2852 enriched genes in NEUROG3-eGFP^+^ cells [36], 277 were bound by NEUROG3, including 56 that were downregulated in the *NEUROG3*^−/−^ cells (Figure 3B-C, Suppl. Table 2). Many of these genes encode for TFs or proteins known to regulate islet cell differentiation and function (see below). Thus, a total of 290 genes specifically expressed in the endocrine lineage (out of 2988) are bound by NEUROG3, suggesting that NEUROG3 directly regulates the expression of about 10% of islet specific genes. In addition, we compared the NEUROG3 cistrome with the human pancreatic adult islet regulome [27]. We found that 782 (90.6%) NEUROG3 binding sites matched with at least one of the adult islet regulatory elements, with 655 (75.90%) of them localized within active enhancers or promoters (Figure 3D and E). This suggests that most of the genes regulated by NEUROG3 are still active in the adult islets, supporting the hypothesis that the transient expression of NEUROG3 at the PEP stage is required to initiate the endocrinogenic program while other transcription factors sustain the transcription of NEUROG3 targets in mature islets by binding to the same regulatory elements.

**Figure 3.**
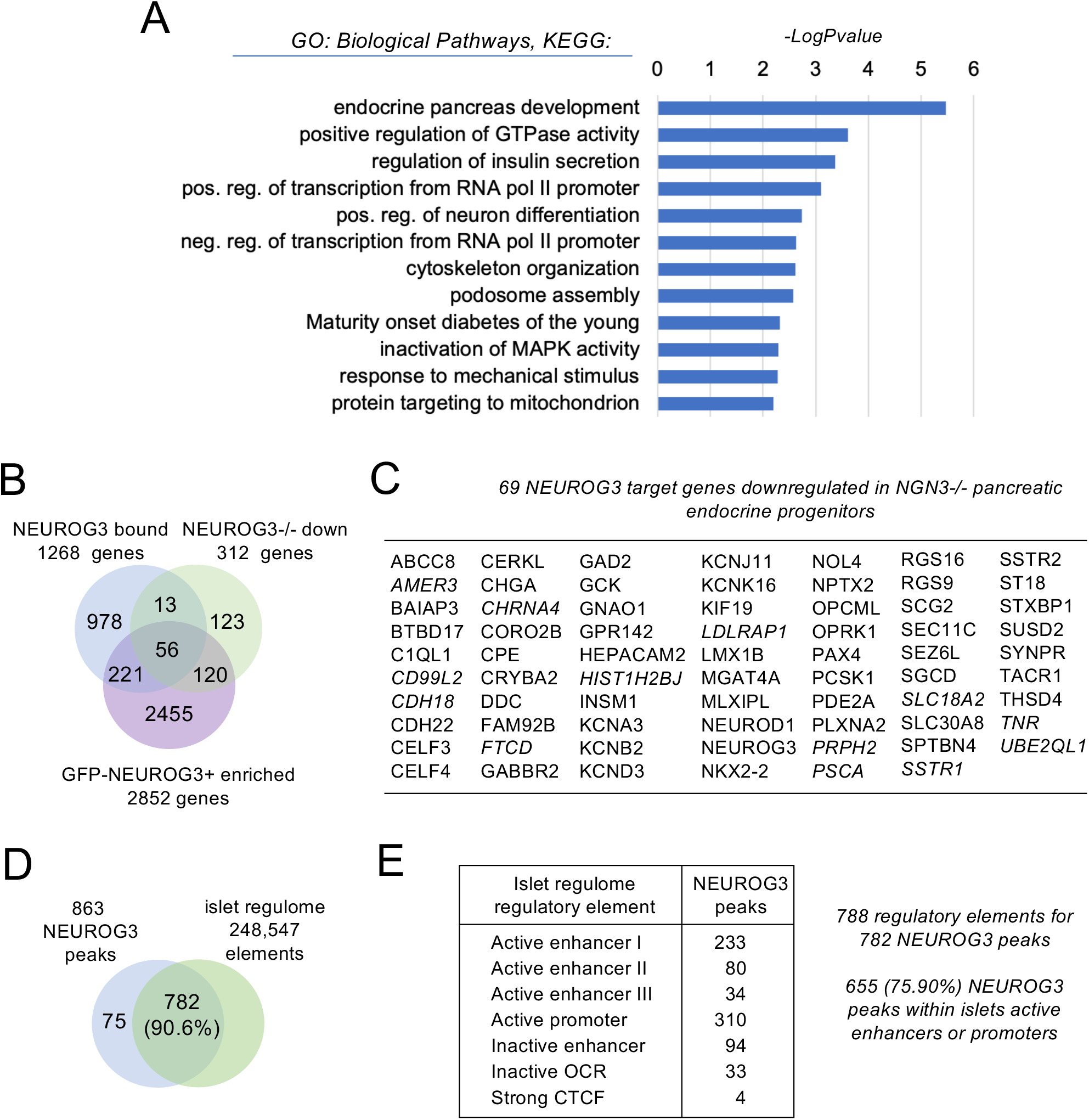
Integration of NEUROG3 occupancy and islet expression of target candidates. (A) Gene ontology analysis (biological process, KEGG pathways) showing selected significantly enriched terms (Log10 p-value ≥2) related to the 1268 NEUROG3 bound genes. (B) Venn diagram illustrating the overlap of NEUROG3 bound genes and the 312 downregulated genes in *NEUROG3*^−/−^ hESC line differentiated to d13 stage (*NEUROG3*^−/−^ down; [34]) and the 2852 genes enriched in NEUROG3-eGFP^+^ hESC line differentiated to pancreatic endocrine progenitors (NEUROG3-eGFP^+^ enriched; [36]). (C) List of the 69 NEUROG3 target genes downregulated in *NEUROG3*^−/−^ PEP. In italics, the 13 bound genes downregulated in *NEUROG3*^−/−^ PEP but not enriched in NEUROG3-GFP^+^ PEP. (D-E) NEUROG3 binding sites matching with one or more islets regulatory element(s) (islet regulome; [27]). OCR, open chromatin regions, CTCF, CTCF binding sites.

### 3.5 NEUROG3 binds to a subset of islet enriched transcription factors

To better understand how NEUROG3 drives islet cell differentiation, we first examined the TF genes bound by NEUROG3. Among the 1268 NEUROG3 bound genes, 138 encode for TFs (Figure 4A and Suppl. Table 2). From those, 24 were enriched in NEUROG3-eGFP^+^ hESC-derived endocrine progenitors, including 8 genes also downregulated in *NEUROG3*^−/−^ cells. Besides the TF genes already mentioned above (*NKX2-2, NEUROD1, NEUROG3, PAX4, INSM1*, and *FOXA2*), we unraveled several other TFs known to control islet cell development in the mouse or human, including *SOX4, RFX3, ST18 (MYT3), MLXIPL, NKX6*.*1* and *LMX1B* (Figure 4A and data not shown), suggesting they could to be regulated directedly by NEUROG3. For instance, NEUROG3 binds to a region in intron 1 of *MLXIPL* (Figure 4B) previously shown to be bound by Rfx6 and Nkx2-2 in the mouse [43; 48]. Interestingly, a NEUROG3 binding site was found 33kb upstream of *SOX4* TSS, and three additional peaks were found within the adjacent *CDKAL1* locus (Figure 4C). The later region is likely to act as a distant enhancer to regulate *SOX4* in islet cells, as suggested by promoter capture HiC data [27] and found to be an activated enhancer (H3K27ac enriched) also at the endocrine progenitor stage [25] (Figure 4C). Thus, while Sox4 has been shown to regulate *Neurog3* expression and be required downstream of *Neurog3* to regulate endocrine differentiation in the mouse [49], *SOX4* might, in turn, be a direct target of NEUROG3. Importantly, we found that NEUROG3 binds to intron 2 of *LMX1B*, a transcription factor recently reported to be critical for generating human islet cells downstream of NEUROG3, suggesting direct transcriptional regulation of *LMX1B* by NEUROG3 (Figure 4D) [25]. Intriguingly, a NEUROG3 peak within the *GLIS3* coding sequence (exon 8) was assigned to both *GLIS3* and *RFX3* (Figure 4E). This peak nicely overlapped with an enhancer region at both endocrine progenitor and adult islets stages [25; 27]. In the adult islets, HiC showed that the two genes are spatially linked [27]. Moreover, RFX3 but not GLIS3 is highly expressed at the endocrine progenitor stage (Figure 4E, [25]) and has recently been documented as a human endocrine fate switch gene regulator [38]. Taken together, these data suggest a possible regulation of *RFX3* by NEUROG3 at the endocrine progenitor stage.

**Figure 4.**
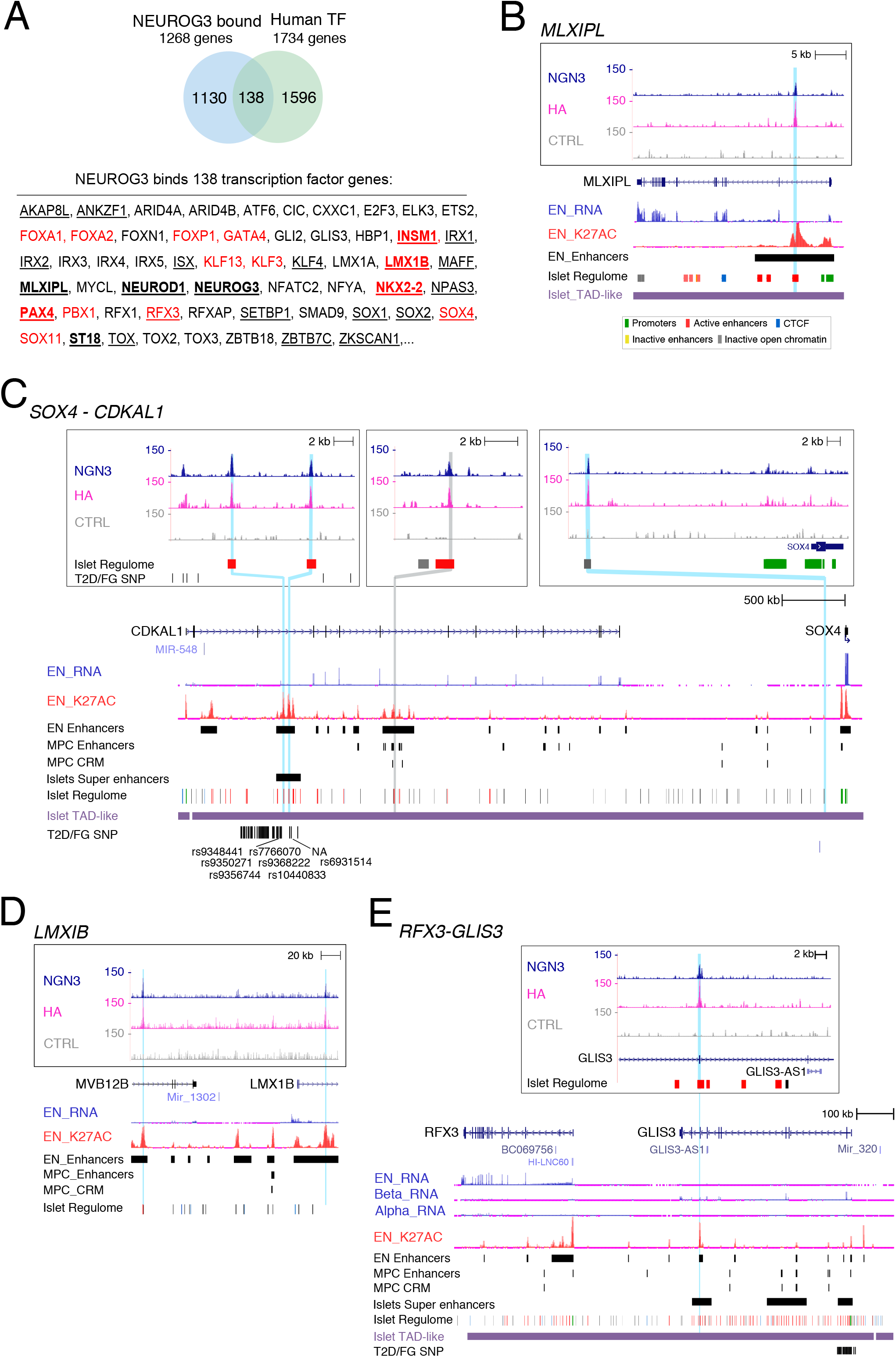
NEUROG3 binding to transcription factors genes. (A) NEUROG3 binds 138 genes encoding human TFs. A selection of TF genes is given. Complete list is given in Suppl Table 2, with the list of 1734 human TFs taken from [32] and https://www.proteinatlas.org. The 24 TF enriched in GFP-NEUROG3+ endocrine progenitors [36], among which 8 are downregulated in *NEUROG3*^−/−^ endocrine progenitors [34] are underlined and in bold, respectively. The 14 TFs belonging to Core Regulatory Circuits (CRCs) in endocrine progenitors as defined by [25] are in red. (B-E) NEUROG3 binding to the *MLXIPL* (B), *SOX4-CDKAL1* (C), *LMXIB* (D) and *RFX3-GLIS3* (E) loci. See Figure 2E for legend description. In (E), RNA-seq data from primary islet beta (Beta-RNA) and alpha (Alpha-RNA) cells are taken from [25].

In a recent study, Alvarez-Dominguez et al. [25] described Core transcriptional Regulatory Circuits (CRCs) for each stages of in vitro beta cells differentiation, based on interconnected autoregulatory loops betweens TFs. Strikingly, out of the 40 TF genes defining the endocrine progenitors CRCs, we show here that 35% are bound by NEUROG3: *LMX1B, FOXA1, FOXA2, FOXP1, GATA4, INSM1, KLF3, KLF13, NKX2-2, RFX3, SOX4, SOX11, PAX4* and *PBX1* (Figure 4A). Of note, since the definition of CRCs relied on TF recognition motifs, NEUROG3, whose motif was not yet known, could not be integrated into the endocrine progenitors CRCs [25]. Importantly, our data reveal and provide molecular mechanistic insights into the role of NEUROG3 as a possible direct regulator of many TFs of the endocrine CRCs.

We further scrutinized the TFs dataset to examine whether NEUROG3 binds to genes known to control islet subtype development and to unveil novel candidates. We focused on transcription factor genes for which NEUROG3 binding site(s) coincided with endocrine progenitor active enhancer regions [25] and were enriched in developing alpha, beta or delta cells based on recent transcriptomic profiling of the human fetal pancreas [37] (Figure 5A). Importantly, an essential role of NEUROG3 in promoting the beta cell fate is supported by its direct regulation of *Pax4* expression, a critical regulator of beta cell development [50]. In addition to *Pax4, Nkx6-1* has been shown to be critical for endocrine progenitors to acquire a beta destiny in the mouse [51]. Supporting a possible direct regulation of *NKX6-1* by NEUROG3, we found a peak at 466kb downstream of *NKX6-1* TSS (Figure 5B). This region overlaps with an endocrine progenitors specific active enhancer region, suggesting that this site might be important for NEUROG3 regulated expression of *NKX6-1* in human islet progenitors. NEUROG3 binding sites were also associated with genes encoding TFs previously reported as markers for beta cells based on their expression, but not yet functionally addressed in endocrine cells development, such as *SMAD9* [52] and *TFCP2L1* [52] (Figure 5C). For *TFCP2L1*, however, the NEUROG3 binding region was not identified as an endocrine progenitor but an adult islet enhancer [27], belonging to an islet-TAD regulating the *GLI2* gene. Whether *TFCP2L1, GLI2*, or both, expressed in human fetal beta cells (Figure 5A), regulate beta destiny in a NEUROG3-dependent manner remains to be established. Of note, we additionally discovered *ETS2* and *ISX* as new NEUROG3 targeted TFs and whose expression is enriched in human feta beta cells, suggesting that they could play a role in human beta cell development (Figures 5A and C).

**Figure 5.**
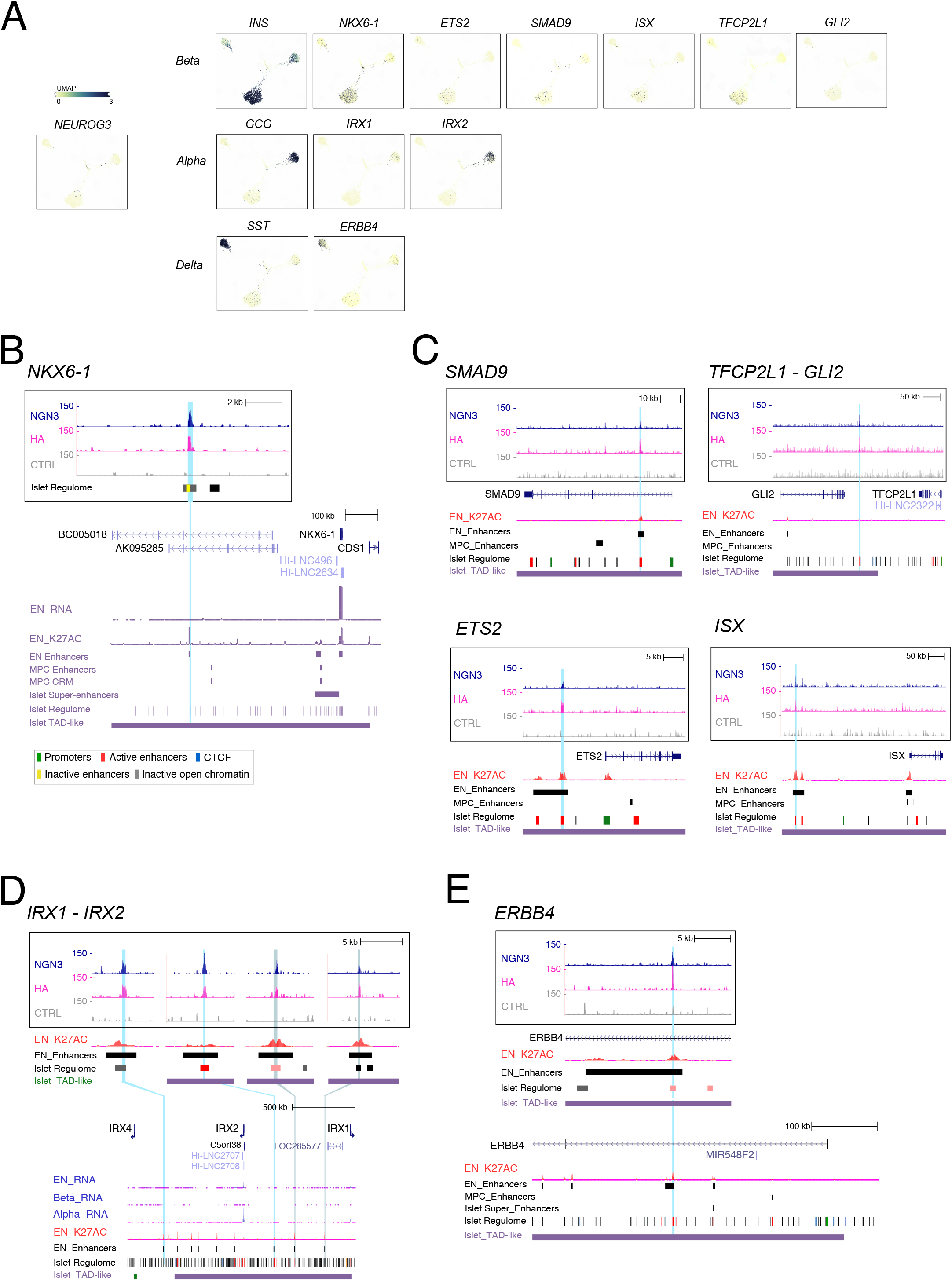
NEUROG3 binds to transcription factors genes enriched in islet-subtypes. (A) Enriched expression of a selection of transcriptional regulators in islet-subtypes (alpha, beta, delta cells) in the human fetal pancreas (taken from https://descartes.brotmanbaty.org). Expression of INS, GCG and SST is provided to define beta, alpha and delta cells, respectively. (B-D) NEUROG3 binding to transcription factor genes expressed in fetal alpha or beta cells: NKX6-1 (B), *SMAD9, TFCP2L1-GL12, ETS2* and *ISX* (C) for beta cells; *IRX1-IRX2* (D) for alpha cells. (E) NEUROG3 binding to the Receptor Tyrosine Kinase/transcription coactivator *ERBB4* gene expressed in developing delta cells. See Figure 2E for legend description.

Regarding alpha destiny, no peaks were assigned to *ARX*, which is essential for alpha cell development in the mouse and human [50; 53]. Of note, NEUROG3 was found bound to regions associated with *IRX1* and *IRX2*, which are both enriched in human fetal (Figure 5A) and adult (Figure 5D and [25; 54]) alpha cells. Interestingly, *Irx2* was induced by ectopic *Neurog3* expression in the chick endoderm [11] and downregulated in hPSC-derived human islet cells lacking ARX [53]. Thus *IRX1/2* are attractive, alpha-specific, NEUROG3 direct targets, although their function in alpha cell development remains to be studied. In contrast to alpha and beta cells, much less is known regarding the regulation of delta cell destiny.

We did not find any binding of NEUROG3 associated with the delta transcription factor HHEX. Nevertheless, our analysis pointed to a possible NEUROG3-dependent candidate regulators of delta cell development. Indeed, we identified a NEUROG3 binding site within the first intron of the EGFR family member Erb-B2 Receptor Tyrosine Kinase 4 (*ERBB4*) gene (Figure 5E), that is highly and specifically expressed in human fetal (Figure 5A) and adult [54] delta cells, and whose ligand neuregulin-4 (NGR-4) was found to be essential for the determination of delta cells in mouse [55]. Of note, ERBB4 is cleaved by gamma-secretase to generate an intracellular domain endowed with TF regulatory activity [56]. Furthermore, during human *in vitro* beta cell differentiation, a gamma-secretase inhibitor is added at the endocrine progenitor stage to inhibit Notch signalling and further promote the beta lineage [39]. Whether the concomitant inhibition of ERBB4, by impeeding the delta destiny, could favor the beta destiny remains to be tested.

Taken together, mapping NEUROG3 occupancy revelaled an unexpectedly broad direct control of TFs in the endocrine gene regulatory network (GRN).

### 3.6 NEUROG3 binds to genes involved in islet cell function

As mentioned above, gene ontology analyses revealed that many NEUROG3 bound genes were associated with insulin secretion, suggesting that NEUROG3 could regulate the expression of genes of the hormone secretory machinary. Indeed, NEUROG3 binding was found in genes linking glucose metabolism to electrical activity in beta cells and subsequent insulin secretion [57], such as the glucose sensor gene *GCK* and *ABCC8/KCNJ11* encoding subunits of the ATP-sensitive K+ channel (Figures 6A-C). Interestingly, other K+ (ATP-independent) channel genes (*KCNA3, KCNB2, KCND3, KCNK16, KCNMA1)*, which also contribute to glucose-stimulated insulin secretion, and are expressed in human fetal islet cells [37], were bound by NEUROG3 (Figure 6A and Suppl. Table 2). In the same line, the voltage-dependent Ca2+ channels (*CACNA1A, CACNA1C, CACNA1E, CACNA2D1, CACNB2*) or genes involved in the formation, composition, or release of secretory granules (*CHGA, SCG2, SLC30A8/ZNT8, SLC18A2/VMAT2, RGS16, RGS4, SYT7, SYT13 SYT3, STX2, STXBP1*) or proinsulin processing (*PCSK1, CPE*) (Figures 6A, D-G, and Suppl. Table 2) are associated to NEUROG3 binding sites. We did not find any binding of NEUROG3 to hormone genes. NEUROG3 binding was also identified in the somatostatin receptor genes *SSTR1, SSTR2*, and *SSTR5* involved in the paracrine regulation of insulin and glucagon secretion [57] (Figure 6H and Suppl. Table 2).

**Figure 6.**
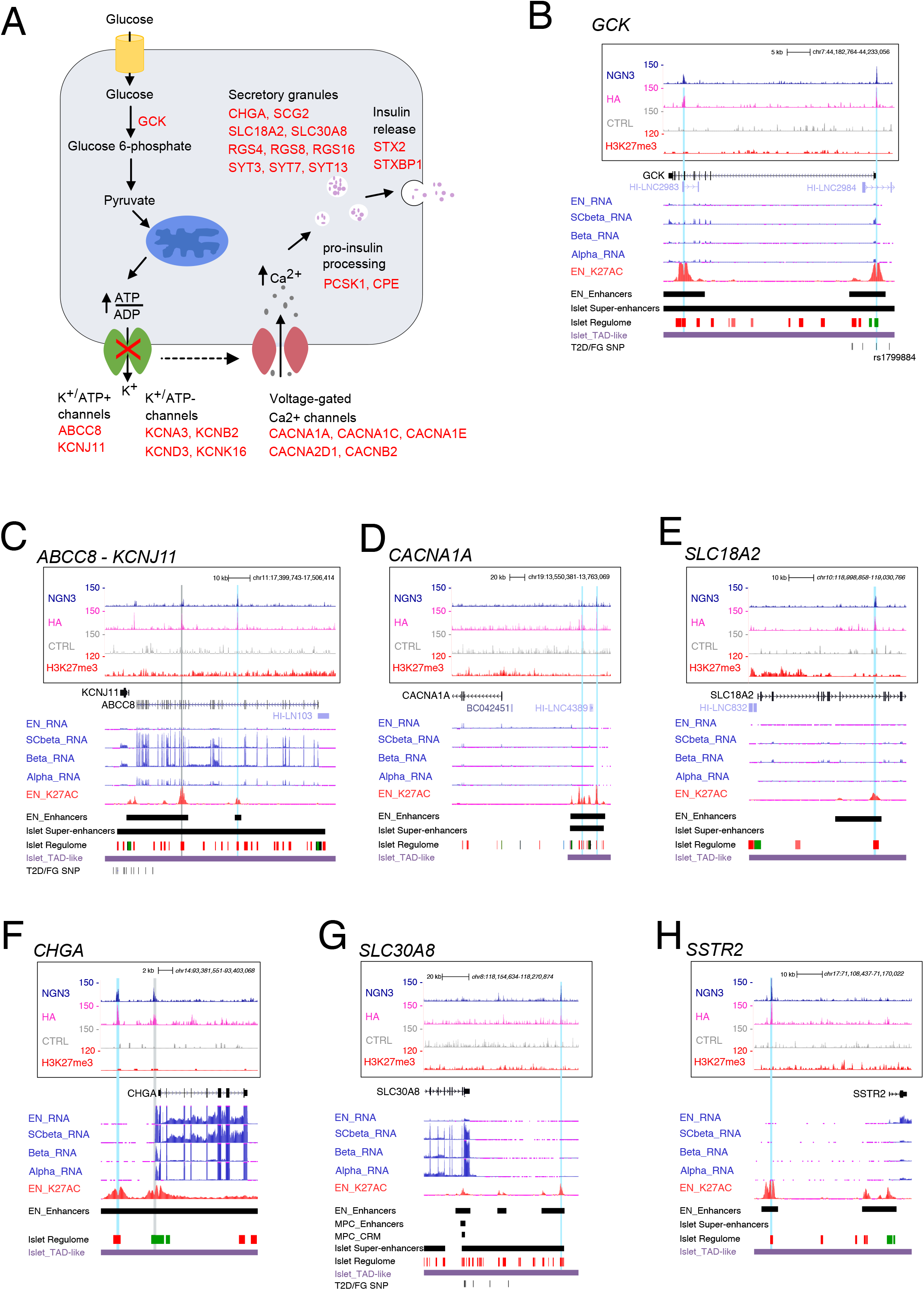
NEUROG3 binding to genes regulating glucose-dependent insulin secretion. (A) Schematic representation of insulin secretion upon glucose sensing in a beta cell. The NEUROG3 bound genes are indicated in red. (B-H) NEUROG3 binding to genes involved in glucose-stimulated insuline secretion: (B) GCK; (C) ABCC8-KCNJ11; (D) CACNA1A; (E) SLC18A2; (F) CHGA; (G) SLC30A8 and (H) SSTR2. See Figure 2E for legend description.

These findings of NEUROG3 bound genes involved in islet cell function were unexpected due to the transient expression of NEUROG3 in endocrine progenitors. Interestingly, several of these target genes, like *ABCC8/KCNJ11, CACNA1A, SLC30A8*, and *SLC18A2*, are not or weakly expressed in endocrine progenitors compared to more differentiated hESC-derived beta (SC-beta) or adult islet cells (Figures 6C, E and G, [25]). We noticed that some of these genes, (e.g., *ABCC8/KCNJ11* and *SLC18A2*, (Figures 6C and E), are decorated with H3K27me3 at or close to their TSS, suggesting that NEUROG3 could prime these genes at the endocrine progenitor stage, but subsequent binding by other TF could be required for their full activation. Thus, NEUROG3 might not only promote islet destiny in uncommitted pancreatic progenitors, but also control the initiation of later generic endocrine programs in maturing islet and beta cells.

### 3.7 NEUROG3 occupancy and ncRNA genes

Non-coding RNAs, such as long non-coding RNA (LnRNA) and microRNA (miRNAs) actively contribute to regulating developmental processes, including pancreatic endocrine specification [33; 58]. We found that 588 (68.3%) NEUROG3 binding sites are less than 100kb distant (98 even within 5kb) from at least one of the human LncRNA expressed in islet cells, compiled by Akerman et al. (2017) [33] (Figure 7A and Suppl. Table 2). Among them, HI-LNC66 (nearby *NEUROD1*), HI-LNC103 (*ABCC8*), HI-LNC4389 (*CACNA1A*), HI-LNC832 (*SLC18A2*) or HI-LNC2984 (*GCG*) could thus as well be targeted by NEUROG3 as their nearby genes (Figures 6B-E and Suppl. Table 2). More generally, this opens the possibility that NEUROG3 could regulate LncRNA expression and consequently their regulatory activity on islet function. Using HOMER annotation, we additionally identified 218 binding sites whose nearest TSS belongs to non-coding genes, among which 66 encode long non coding intergenic LincRNAs, 31 miRNA, and 23 antisense RNA (Figure 7A and Suppl. Table 2). Several ncRNA, not described in Akerman et al. (2017) list [33], were antisense to genes already mentioned above as bound and therefore possibly regulated by NEUROG3, such as *GLIS3, ISX, KCND3, KCNMA1*, and *SSTR5*. NEUROG3 could therefore regulate these genes directly or indirectly by regulating the expression of their antisense RNA. In line with this hypothesis, the somatostatin receptor SSTR5-AS1 is downregulated in *NEUROG3*^−/−^ PEP cells, whereas SSTR5 is enriched in NEUROG3-eGFP^+^ PEP cells (Suppl. Table 2). *LINC00261*, nearby to *FOXA2*, is highly expressed during pancreatic differentiation [25; 58] (Figure 2E) and was shown recently to be required for the efficient differentiation of hESCs to insulin-producing cells [58], through the trans-regulation of *PAX4* and *MAFB* TFs, whereas its *cis*-regulatory role on *FOXA2* is debated [58; 59]. *FOXA2* but not *LINC00261* is downregulated in *NEUROG3*^−/−^ PEP (Suppl Table 2). Therefore, whether NEUROG3 regulates *LINC00261*, or *FOXA2*, or both, is an open question that can be extended to other NEUROG3 bound genes with ncRNAs in their vicinity.

**Figure 7.**
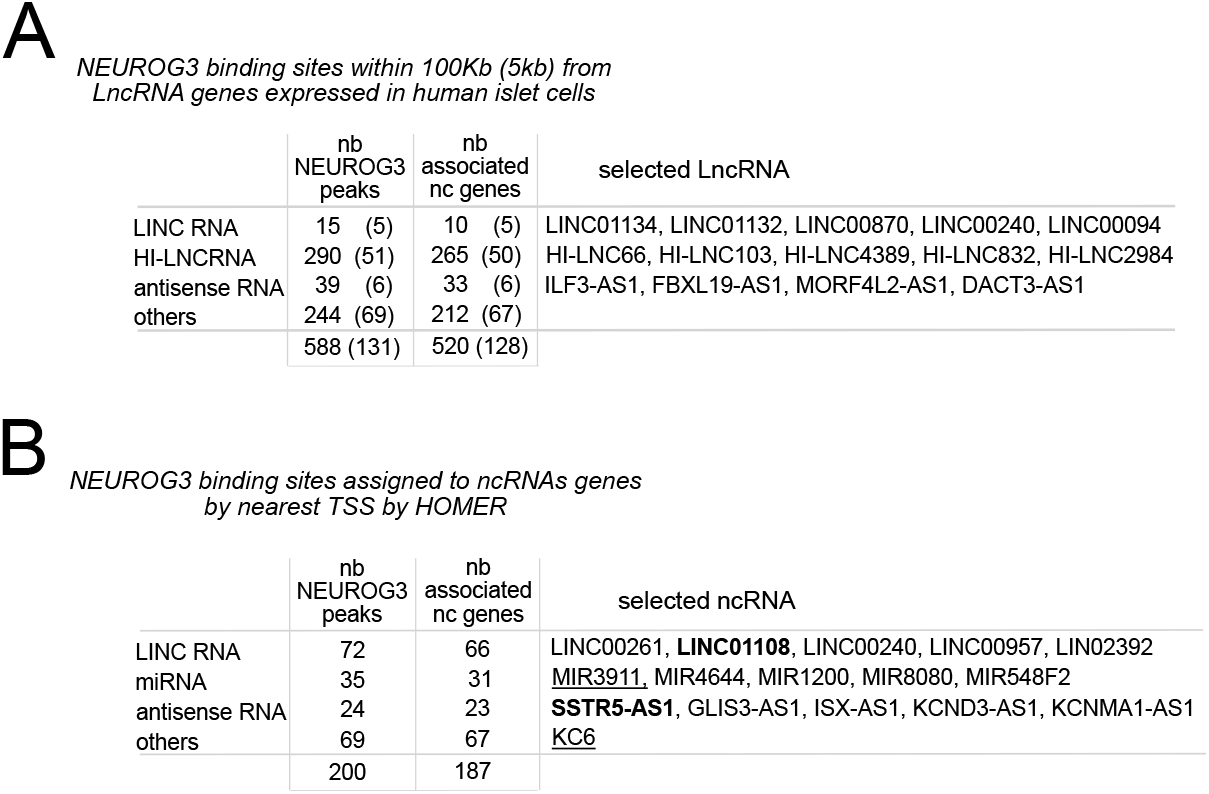
NEUROG3 binds to ncRNA genes. (A) Distribution of NEUROG3 binding sites and number of associated non coding (nc) genes located within 100kb (or 5kb from the TSS) of human beta-cell enriched lncRNAs, taken from [33]. HI-LNC, human islet long non coding RNA. (B) Distribution of NEUROG3 binding sites within ncRNA genes, including LINC RNA, miRNA, antisense RNA and other RNA annotated by HOMER. Some selected examples are given, underlined when found enriched in NEUROG3-eGFP^+^ PEP and in bold when downregulated in *NEUROG3*^−/−^ PEP.

### 3.8 NEUROG3 binding at T2DM risk variants

Genome-wide association studies (GWAS) have identified hundreds of genetic variants associated with increased T2DM susceptibility [60]. It is essential to understand how these T2DM linked SNPs contribute to the disease, which genes they affect and how, whether it is by altering the protein sequence or, most frequently, distal cis-regulatory elements. Miguel-Escalada et al. [27] have compiled a list of 23,154 genetic variants associated with T2D and/or fasting glycemia (T2D/FG SNPs) within 109 loci. We found an overlap between 7 risk loci (harboring 152 SNPs) and NEUROG3 bound endocrine progenitor enhancers (Figures 8A-B and Suppl. Table 2). Moreover, 4 of these risk loci had additional SNPs, not in the abovementioned endocrine enhancers but falling within a NEUROG3-binding site (Figures 8A-B and Suppl. Table 2). A closer examination of the genomic regions of some of these 7 risk loci was performed. The risk alleles rs1799884, rs635299, and rs114152784 lie within NEUROG3-binding sites at the promoter regions of *GCK, SIX5*, and *MDC1*, respectively (Figures 6B, 8C and data not shown) and rs7245708 at the bi-directional promoter of QPCTL and SNRPD2 (Figure 8C). *SIX5, QPCTL*, and *SNRPD2* belong to the GIPR GWAS locus, and whereas all 4 genes are variably expressed at the endocrine progenitor stage (Figure 8C and [25]), GIPR is the most highly expressed in the human fetal pancreas [37]. The NEUROG3 binding sites within the *CDKAL1* locus coincide with several T2D-FG SNPs previously assigned to the distal gene *SOX4* (Figure 4C and [27]). The UBE2Z risk region contains several genes expressed at the endocrine progenitor stage and in the fetal pancreas that could be affected by SNPs, such as *ATP5G1, CALCOCO2, TTLL6*, and *UBE2Z* itself (Figure 8D and [37]). A NEUROG3 binding site lies upstream TTLL6 TSS (−1255bp) that is expressed higher at the endocrine progenitor stage than in beta cells (Figure 8D, [25]). In mouse, *TTLL6* expression follows that of NEUROG3 [61] and mouse invalidated for the *Ttll6* gene revealed decreased circulating glucose level (https://www.mousephenotype.org/data/genes/MGI:2683461). Altogether, these data support the potential regulation of *TTLL6* by NEUROG3. Interestingly, the NEUROG3 binding site assigned to the *ARAP1* gene coincided with an islet active enhancer (R11, Figure 8E) belonging to a regulatory region of the *STARD10* and *FCHSD2* genes, highly enriched in T2D/FG variants [27; 62]. The variable region (VR) within this locus was demonstrated to be required for insulin secretion [62]. *STARD10* is already expressed at the endocrine progenitor stage and in the fetal pancreas (Figure 8E and [25; 37]), but whether this expression is regulated by NEUROG3 and this regulation affected by genetic variants remain to be established.

**Figure 8.**
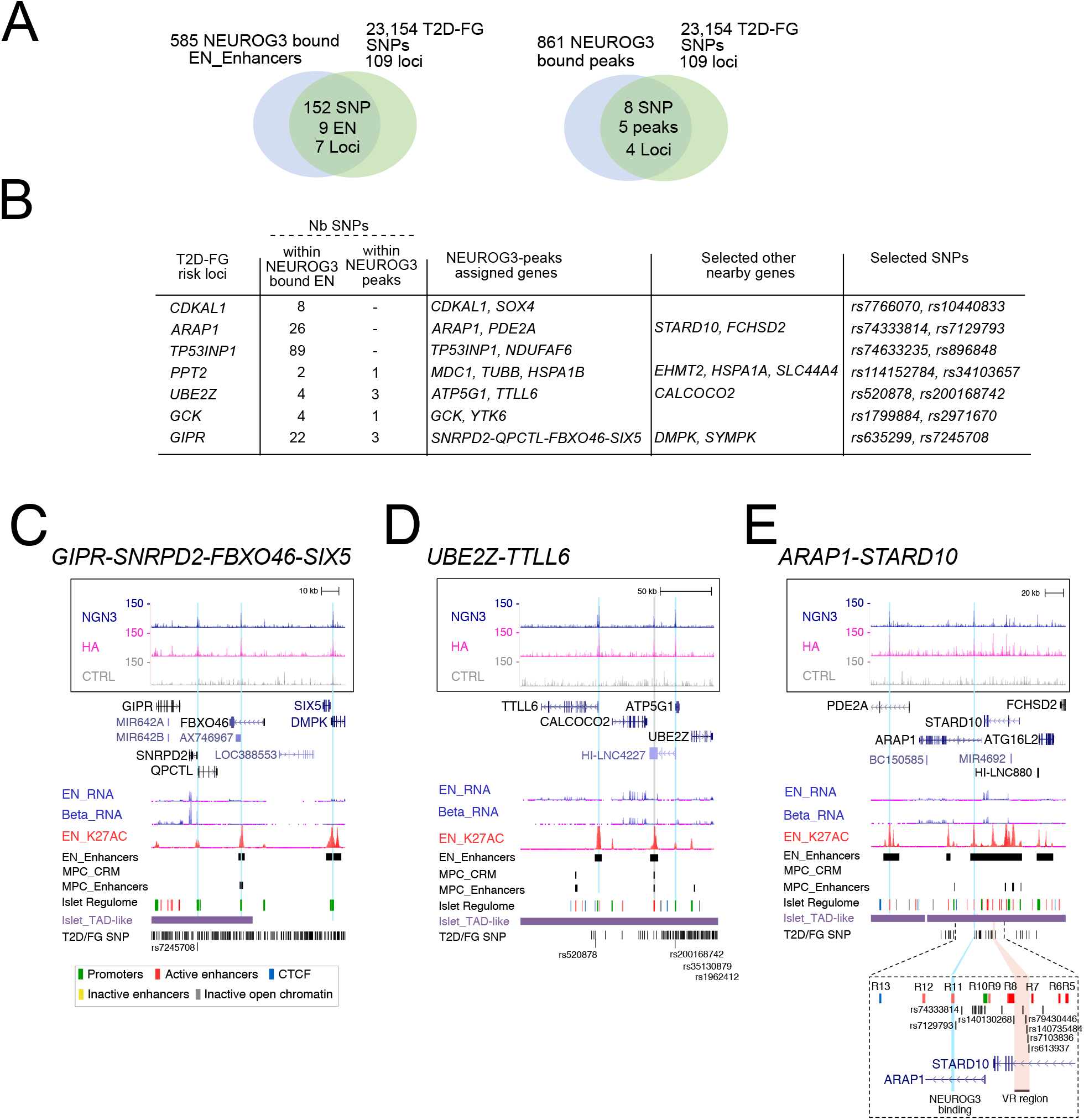
T2D and fasting glycemia (FG)-associated genetic variants located within NEUROG3-bound EN enhancers and NEUROG3-bound regions. (A) Venn diagram illustrating the overlap of NEUROG3 bound EN enhancers (left, [25]) or peaks (right) and the 23,154 T2D-FG SNPs distributed over 109 risk loci compiled by [27]. (B) The 7 risk loci overlapping with NEUROG3-bound enhancers or peaks, with the number of SNPs, NEUROG3-peaks assigned genes, some selected nearby genes and selected SNPs given for each risk loci. (C-E) NEUROG3 binding to the *GIPR-SNRPD2-FBXO46-SIX5* (C), *UBE2Z-TTLL6* (D) and *ARAP1-STARD10* (E) loci. In E, the inset is a magnification of the region previously described as a regulatory region of the *STARD10* and *FCHSD2* genes [27; 62]. R5-13, regulatory regions; VR, variable region. See Figure 2E for legend description.

Taken together, the overlap of T2DM SNPs and NEUROG3 bound regions suggests that T2DM susceptibility could arise from impaired, NEUROG3-dependent genetic programs.

## 4. CONLUSION

Despite the major progress made in the generation of functional beta cells from pluripotent stem cells for a cell therapy in diabetes, directed differentiation protocols lack robustness, and obtaining glucose-responsive cells remains difficult. The overall strategy was to mimic pancreas and islet developmental programs identified essentially in rodents. While the successful production of insulin-producing cells from PSC *in vitro* attests that these programs are remarkably conserved, it is important to acquire additional insights into the gene regulatory networks controlling islet cell development in human to optimize differentiation protocols. Notwithstanding the essential function of NEUROG3 in islet cell development in mouse and human, its downstream direct targets implementing the endocrinogenic program are essentially unknown. Searching and analyzing NEUROG3 binding sites in purified hiPSC-derived PEP, using the CUT&RUN technique, revelead more than a thousand novel putative direct targets. Importantly, NEUROG3 binding largely overlaps with PEP active enhancers (H3K27ac binding) as defined by others [25], underlining the importance of NEUROG3 in promoting gene expression in PEPs. Our study revealed that NEUROG3 binds to a high number of important islet TFs as well as novel possible transcriptional regulators of islet cell differentiation. Moreover, a plethora of genes involved at several key steps of the insulin secretion pathway is bound by NEUROG3. In addition, we unveiled a panel of ncRNA potentially regulated by NEUROG3. Finally, we reveal that NEUROG3 binding overlaps with a series of T2DM associated SNPs. Taken together, our results suggest that NEUROG3 controls the progression of islet cell differentiation as well as the setting up of the hormone secretory machinery. The pleiotropic functions of NEUROG3 direct targets support the severity of NEUROG3 mutations in mice and humans as well as the potential of NEUROG3 to induce an endocrinogenic program when expressed ectopically. To our knowledge, this is the first genome wide characterization of NEUROG3 occupancy in iPSC-derived PEPs.

## AUTHOR CONTRIBUTION

V.S., R.M., E.G.S, A.K., A.M. and S.G. performed iPSC gene editing, differentiations and characterizations. V.S. and R.M. performed the CUT&RUN experiments, B.J. the Illumina sequencing and T.Y, V.S and S.J. the bioinformatics analyses. C.B. produced the pA-MN. C.H. provided the SB AD3.1 line and expertise for iPSC culture. K.H.L and P.S. performed the RNA-seq data for NEUROG3^***−/−***^ iPSC line and participated in the manuscript redaction. V.S. and G.G. conceived the work, analyzed the data and wrote the manuscript. G.G. obtained financial support.

## ACKNOWLEDGMENTS

The authors thank the members of the Gradwohl team and the Genomeast platform (particularly Christelle Thibault-Carpentier and David Rodriguez), Flow cytometry and Cell culture facilities for the sequencing of the CUT&RUN samples, cell sorting and hiPSC maintenance respectively. The authors are grateful to I. Cebola for providing ChIP-seq data and R. Scharfmann for helpful discussions. The Gradwohl lab is funded by the Novo Nordisk Foundation (Challenge Grant NNF14OC0013655). Sequencing was performed by the GenomEast platform, a member of the ‘France Génomique’ consortium (ANR-10-INBS-0009). This work used the Integrated Structural Biology platform of the Strasbourg Instruct-ERIC center IGBMC-CBI supported by FRISBI (ANR-10-INBS-0005-001). IGBMC is supported by the grant ANR-10-LABX-0030-INRT, a French State fund managed by the Agence Nationale de la Recherche under the frame program Investissements d’Avenir ANR-10-IDEX-0002-02.

## APPENDIX A. SUPPLEMENTARY DATA

**Supplemental Figure 1.**
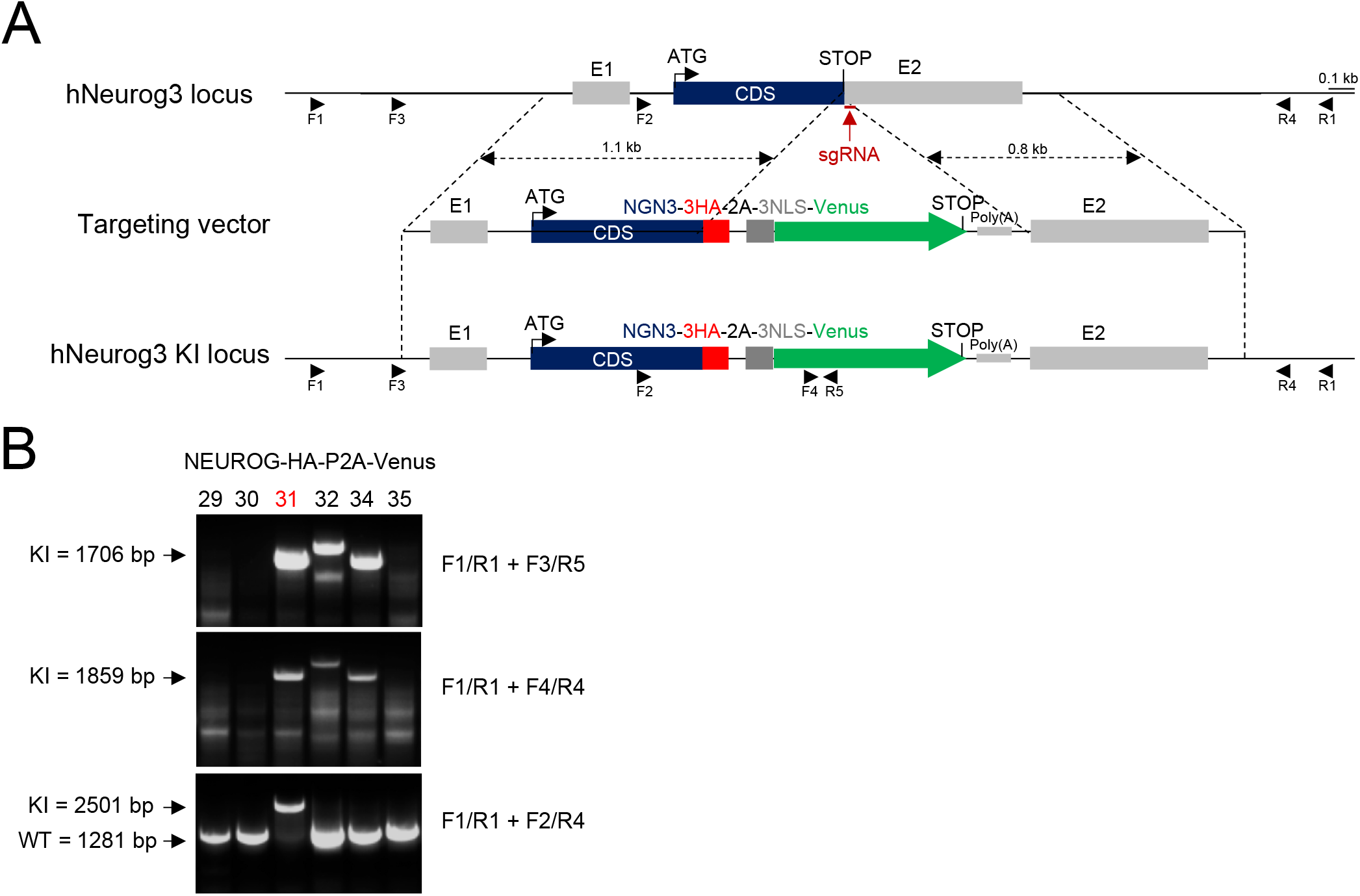
Generation of the NEUROG3-HA-P2A-Venus hiPSC line by CRISPR/Cas9. (A) CRISPR/Cas9 mediated targeting of the NEUROG3 locus to knock in the 3HA-P2A-3NLS-Venus cassette in fusion with NEUROG3 coding sequence. Position of the sgRNA and the primers used for genotyping and sequencing is indicated. (B). Genotyping by PCR using the indicated primers. The clone used throughout this study is clone#31. See Suppl. Table 1 for oligonucleotide sequences.

**Supplemental Figure 2.**
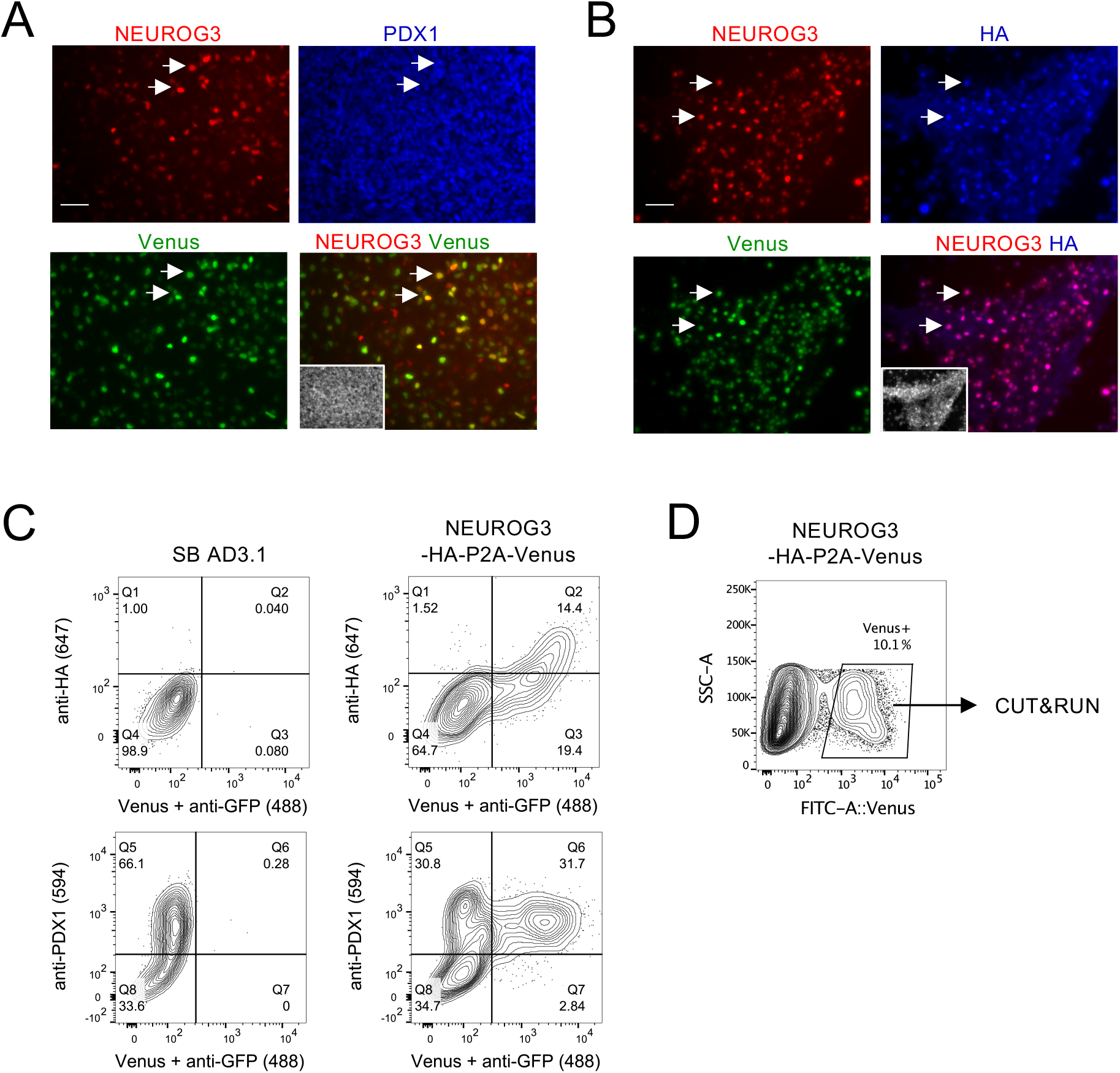
Differentiation of NEUROG3-HA-P2A-Venus hiPSCs line to pancreatic endocrine progenitors (PEP) and FACS-sorting of Venus+ cells. (A-B) Immunofluorescence staining for NEUROG3, Venus and PDX1 (A) or HA (B) at PEP stage (day 13 of differentiation). DNA counterstained with Dapi is shown as an inset within the merged images. Scale bar = 50 µM. Arrows point to a selection of cells co-expressing NEUROG3, Venus and PDX1 (A) or NEUROG3, Venus and HA (B). (C) Representative flow cytometry plots showing the percentage of cells expressing NEUROG3-3HA (anti-HA antibody), Venus (anti-GFP anibody) and PDX1 (anti-PDX1 antibody) at day 13 for the differentiated SB AD3.1 and NEUROG3-HA-P2A-Venus hiPSC lines. (D) Representative flow cytometry plot showing the sorted living Venus+ cells at day 13 used for CUT&RUN experiments.

**Supplemental Figure 3.**
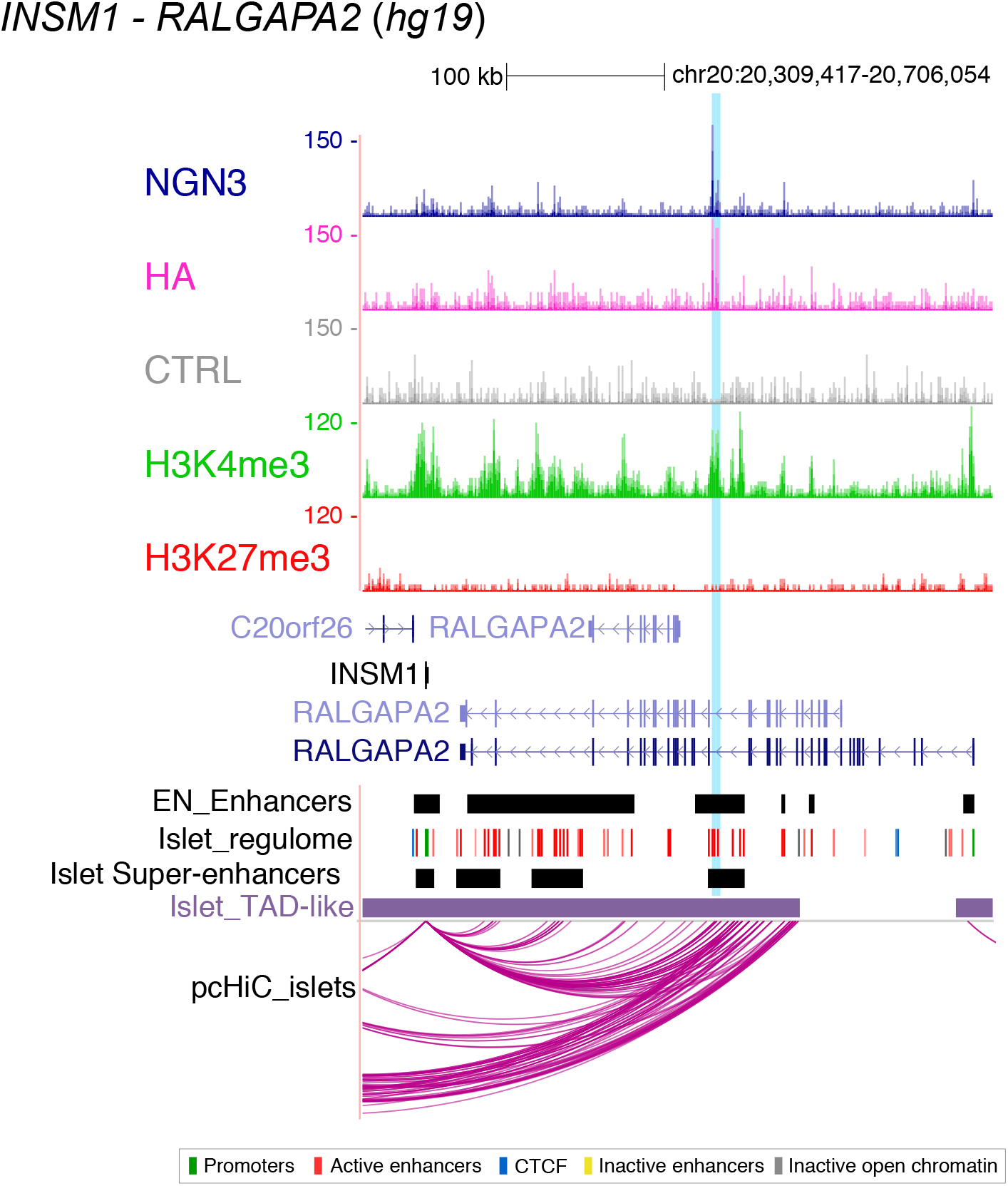
NEUROG3 binds to *INSM1-RALGAPA2* locus. Genome browser tracks showing NEUROG3, HA, H3K4me3, H3K27me3 and the CTRL CUT&RUN data at the *INSM1-RALGAPA2* locus. Coordinates are from hg19. The position of NEUROG3 binding sites is highlighted in light blue. Position of endocrine progenitor enhancers (EN Enhancers) were taken from [25]. Data from adult islets (Super-enhancers, Islet regulome, TAD-like regions and promoter capture HiC pc-HiC_islets) are taken from [27], isletregulome.org and http://epigenomegateway.wustl.edu/.

**Supplemental Figure 4.**
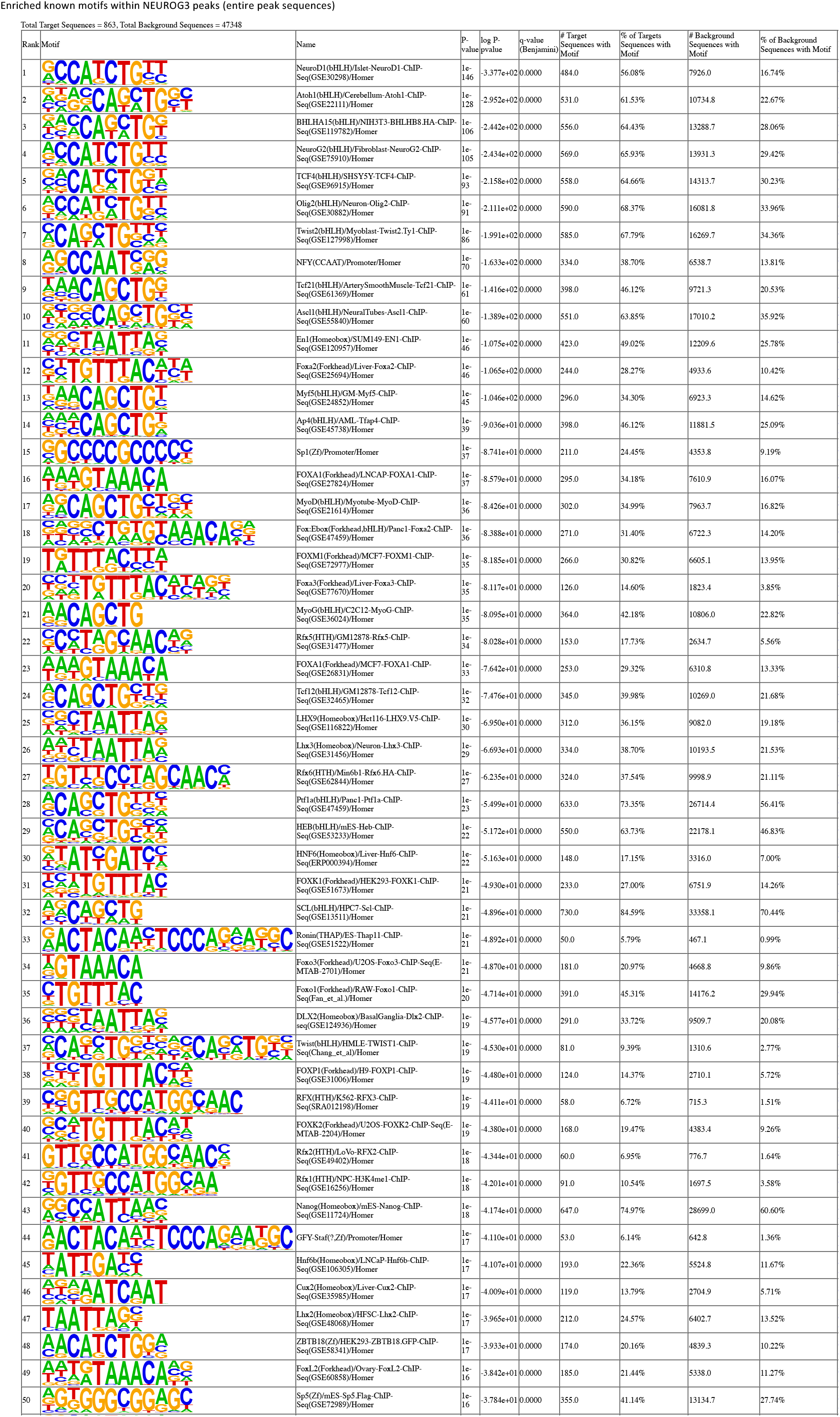
**List of the 50 most significantly enriched TFs known motifs in NEUROG3 binding sites,** in regions defined by the entire peak coordinates.

**Supplemental Table 1**.

ST1.1. List of oligonucleotides

ST1.2. List of antibodies

ST1.3. hiPSC differentiation protocol and media

**Supplemental Table 2**.

ST2.1. The 863 NEUROG3 binding sites, identified by CUT&RUN with both the anti NEUROG3 and anti HA antibodies, and annotated with HOMER v3.4

ST2.2. The 863 NEUROG3 binding sites are assigned to 1268 unique genes by GREAT v4.0.4

ST2.3. Co-occurence of NEUROG3, FOXA2 and RFX6 motifs in the NEUROG3 peaks

ST2.4. Gene ontology performed with https://DAVID.org on the 1268 NEUROG3 bound genes

ST2.5. Genes downregulated in NEUROG3^−/−^ hESC line differentiate to pancreatic endocrine progenitors

ST2.6. NEUROG3 bound genes and expression in islet lineage

ST2.7. NEUROG3 bound TFs genes

ST2.8. T2D and FG-associated genetic variants located within NEUROG3-bound EN enhancers and/or NEUROG3-bound regions.

